# Early blindness shifts the feedforward laminar profile of hMT+/V5 from visual to auditory motion

**DOI:** 10.64898/2026.09.04.749307

**Authors:** Marco Barilari, Laurentius (Renzo) Huber, Jacek Matuszewski, Eléonore Giraudet, Remi Gau, Marc Van Baelen, Olivier Collignon

## Abstract

The occipital cortex of people born blind massively enhances its response to sounds, but the brain circuitry supporting such crossmodal plasticity remains elusive. Here we capitalized on ultra-high-field fMRI (7T) coupled with sub-millimetre BOLD and VASO acquisitions to infer whether motion-related information is processed via feedforward or feedback pathways, probing responses to visual motion in sighted participants and auditory motion in both sighted and early blind individuals. By identifying circuitry from layer-dependent activity of the middle temporal cortex (hMT+/V5), we observed that moving, but not static, sounds selectively elicited a feedforward response in the middle layers of hMT+/V5 of blind people, analogous to visual motion processing in sighted individuals. Furthermore, we observed that hMT+/V5 shows enhanced auditory motion-selective connectivity with the Planum Temporale in the sighted and with the cuneus in blind people. These findings reveal that hMT+/V5 implements a feedforward motion selective response profile in sighted and blind individuals, with its sensory input shifting from vision to audition in the absence of visual experience.

## INTRODUCTION

The study of how the visual system adapts to early blindness (EB) has long served as a leading model to investigate how experience influences genetic programs of brain development (Bavelier & Neville, 2002; Frasnelli et al., 2011).The enhanced response to sounds or touch observed in the occipital cortex of early blind humans (Cohen et al., 1997; Sadato et al., 1996) and other animals (Kahn & Krubitzer, 2002; Yaka et al., 2000) is arguably one of the most striking demonstrations of experience-dependent brain plasticity.

While crossmodal responses in the occipital cortex of early blind humans have been extensively characterized using standard functional Magnetic Resonance Imaging (fMRI) at the macroscale, these phenomena remain poorly understood at the circuit (mesoscale) level. Recent advances in ultra-high-field laminar fMRI now allow non-invasive measurement of cortical depth-resolved activity, enabling inferences about feedforward vs. feedback input based on established laminar hierarchies (Lawrence et al., 2019). Invasive work in animals shows that primary sensory inputs arrive predominantly in middle layers (feedforward), whereas non-primary or contextual signals arise in superficial and deep layers (feedback) (Oude Lohuis et al., 2024; Schroeder & Foxe, 2002). A growing number of laminar fMRI studies in humans corroborate these principles, showing that visual input in the occipital cortex mainly activates middle layers while predictive, contextual or multisensory information mostly engages superficial and/or deep cortical layers (Aitken et al., n.d.; Chai et al., 2021; Dresbach et al., 2023; Gau et al., 2020; Huber et al., 2015; Lankinen et al., 2022; Lawrence et al., 2019; Muckli et al., 2015). However, the laminar expression of crossmodal responses in the occipital cortex of blind people has never been investigated.

Characterizing the laminar profile of responses to visual and auditory motion in the middle temporal cortex of sighted and blind would provide new insights into the development and plasticity of multisensory brain circuits due to sensory experience. Indeed, it has been shown that hMT+/V5, a region known for being selective to visual (Watson et al., 1993; Zeki, 1991) and to a lesser extent auditory motion in sighted people (Poirier et al., 2006; Rezk et al., 2020), robustly enhances its functional preference to moving sounds in early blind people (Battal, 2018; Dormal et al., 2016; Poirier et al., 2006). The preferential responses evoked by moving sounds in the hMT+/V5 of blind people match the magnitude, spatial position, and functional selectivity of visual motion responses in sighted controls (Battal et al., 2019; Dormal et al., 2016; Poirier et al., 2006). Despite decades of research, the pathways by which auditory motion signals access those middle temporal regions in blindness remain unknown. The main proposed mechanistic explanations include either subcortical rerouting or strengthened cortico-cortical feedback from auditory motion-selective regions (Bavelier & Neville, 2002; Bronchti et al., 2002; Dormal et al., 2012; Karlen et al., 2006).

Does non-visual information access “visual” cortex by reorganizing early feedforward thalamo-cortical sensory pathways or, alternatively, does it rely on scaling up cortico-cortical feedback mechanisms presumably existing in sighted individuals? This distinction is fundamental: feedforward access would indicate that auditory signals have been incorporated into canonical motion-processing circuits similar to the one observed for vision in sighted, whereas feedback-driven activation would suggest that auditory motion is first computed in other cortical regions before reaching hMT+/V5 via top-down influences (Felleman & Van Essen, 1991; Macaluso et al., 2000; Macaluso & Driver, 2005).

Here, we address this question by testing Sighted and Early Blind individuals using ultra-high-field (7T) MRI combining an independent whole brain functional localizer and depth-resolved high-resolution functional imaging acquired over a restricted field of view (Fig.1a-c). This approach allowed us to establish the laminar signatures of visual and auditory motion responses in hMT+/V5 in sighted observers, and then test how this circuitry is affected by early blindness. Crucially, to achieve laminar specificity, we relied on cerebral blood– volume–weighted Vascular Space Occupancy (slice-saturation slab-inversion or SS-SI VASO) imaging (Huber et al., 2014), which is less sensitive to signal contamination from intracortical and pial draining veins than conventional gradient-echo BOLD fMRI. Whereas laminar BOLD responses are strongly biased toward superficial layers due to venous drainage, VASO preferentially reflects microvascular changes more closely linked to local neural activity, providing a more faithful estimate of depth-dependent input organization (Finn et al., 2019; Huber et al., 2017; Kim, 2025). By leveraging the complementary strengths of VASO and BOLD imaging, our approach enables a robust dissociation of feedforward and feedback signals within hMT+/V5 and offers a principled framework to probe circuit-level mechanisms of crossmodal plasticity in the human brain.

**Figure 1.**
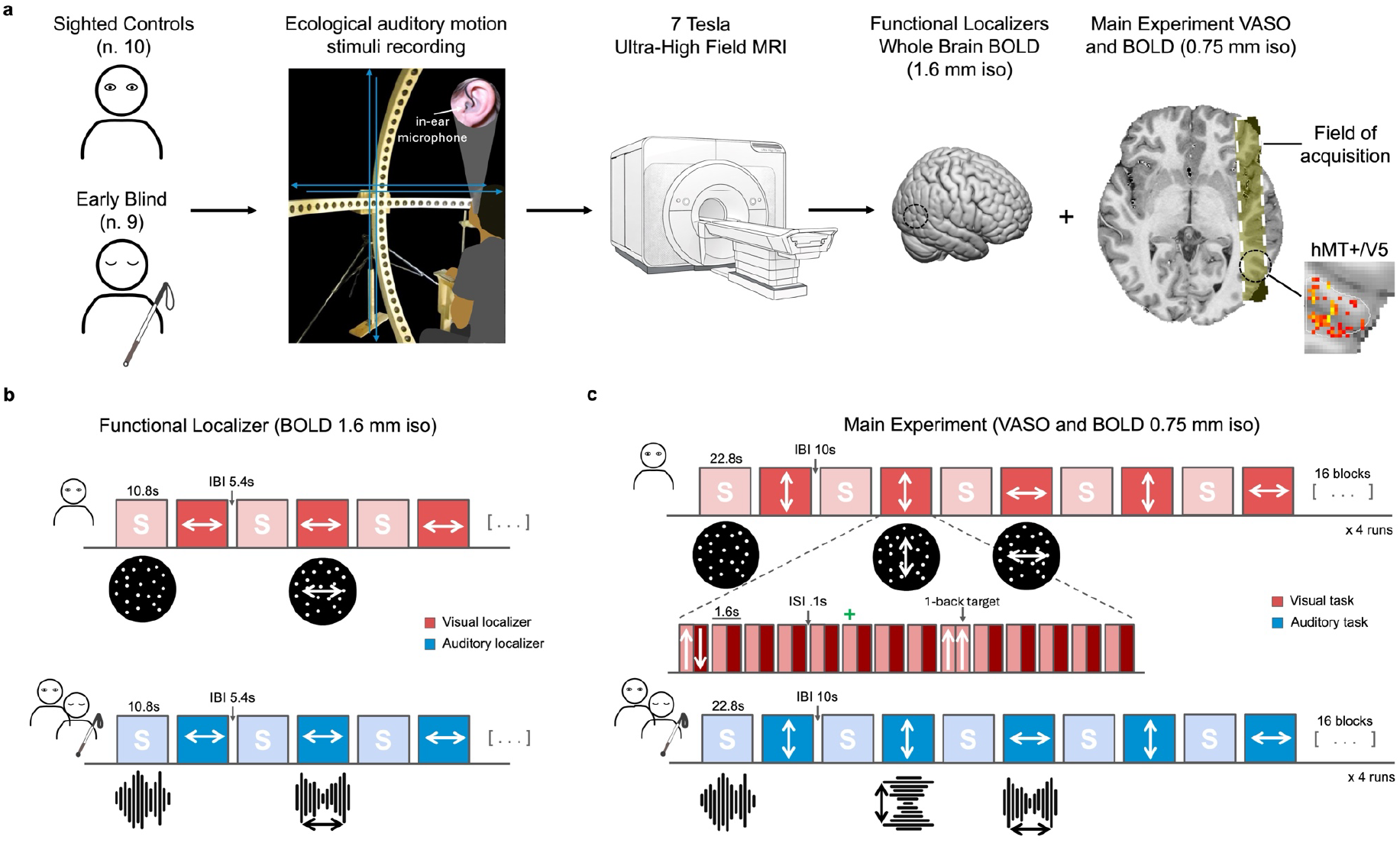
Experimental design and acquisition strategy. a, Overview of the data acquisition workflow in Sighted Controls (SC) and Early Blind (EB) participants. For each participant, ecologically valid auditory motion stimuli were generated individually by recording sounds using in-ear binaural microphones while participants (represented as a silhouette) were seated at the centre of a custom speaker array composed of two semicircular arms arranged in the horizontal and vertical planes. This procedure preserved individual head- and pinna-related acoustic filtering. The recorded sounds were subsequently replayed during 7-Tesla fMRI acquisition for the auditory tasks. All participants underwent (i) whole-brain blood-oxygen-level-dependent (BOLD) imaging for the visual and auditory localizers (1.6 mm isotropic resolution), and (ii) submillimetre slice-saturation slab-inversion Vascular Space Occupancy (SS-SI VASO) and BOLD acquisition (0.75 mm isotropic resolution). To meet the spatial resolution requirements of laminar fMRI, data were acquired using a restricted field of view. The acquisition slab was positioned sagittally over the right hemisphere to optimally cover hMT+/V5 in each participant based on anatomical landmarks. **b,** Schematic of the visual and auditory motion localizer runs, each consisting of alternating blocks of static and moving stimuli (horizontal axis) presented in the corresponding sensory modality. **c,** Design of the main experiment acquired using VASO sequence at 0.75 mm isotropic resolution. Four runs were acquired per modality. Runs consisted of alternating static and moving blocks. Motion stimuli were presented along either the horizontal (left-right) or vertical (up-down) axis. Visual motion consisted of white dots presented on a black screen while auditory motion consisted of individually recorded moving pink noise stimuli generated prior to MRI acquisition.

## RESULTS

### Visual and auditory motion selectivity in sighted and blind individuals

First, to individually localize motion-selective region of interest (fROI) for subsequent laminar analyses, participants underwent whole-brain fMRI at 1.6-mm isotropic resolution while viewing (sighted) or listening (sighted and blind) to static and moving stimuli (Fig.1b). We computed a motion-greater-than-static contrast [motion>static] which identified subject-specific functional ROIs in occipito-temporal cortex: in sighted controls (SC), a visually defined hMT+/V5 and an auditory-responsive hMTa; in early blind (EB) individuals, an auditory-defined hMT+/V5. Across individuals, motion selectivity consistently emerged near the junction of the lateral occipital and the ascending limb of the inferior temporal sulcus, the canonical location of hMT+/V5 (Dumoulin, 2000) (Fig2b,d,e).

At the group level, visual motion compared to static preferentially activated bilateral hMT+/V5 and V1 in SC (Fig.2a left). Auditory motion contrasted to static sounds elicited in both groups a preferential response in the temporal cortex (in particular in the Planum Temporale) as well as in middle temporal regions in EB (Fig.2a centre), and to a lesser extend in sighted people (Fig 2a left). These motion selective responses in middle temporal region for sound and sight were clearly observable in single participants as well (Fig.2e) and partially overlapped (Fig. 2b). When contrasting auditory motion selective response (greater than static) between sighted and blind, we observed enhanced auditory motion selectivity in bilateral hMT+/V5 and bilateral Cuneus of the blind (Fig.2a right, Table S2). Unsmoothed beta estimates from the individually defined ROIs (shown for visualization only) illustrate the motion selective response of those regions (Fig.2c).

**Figure 2.**
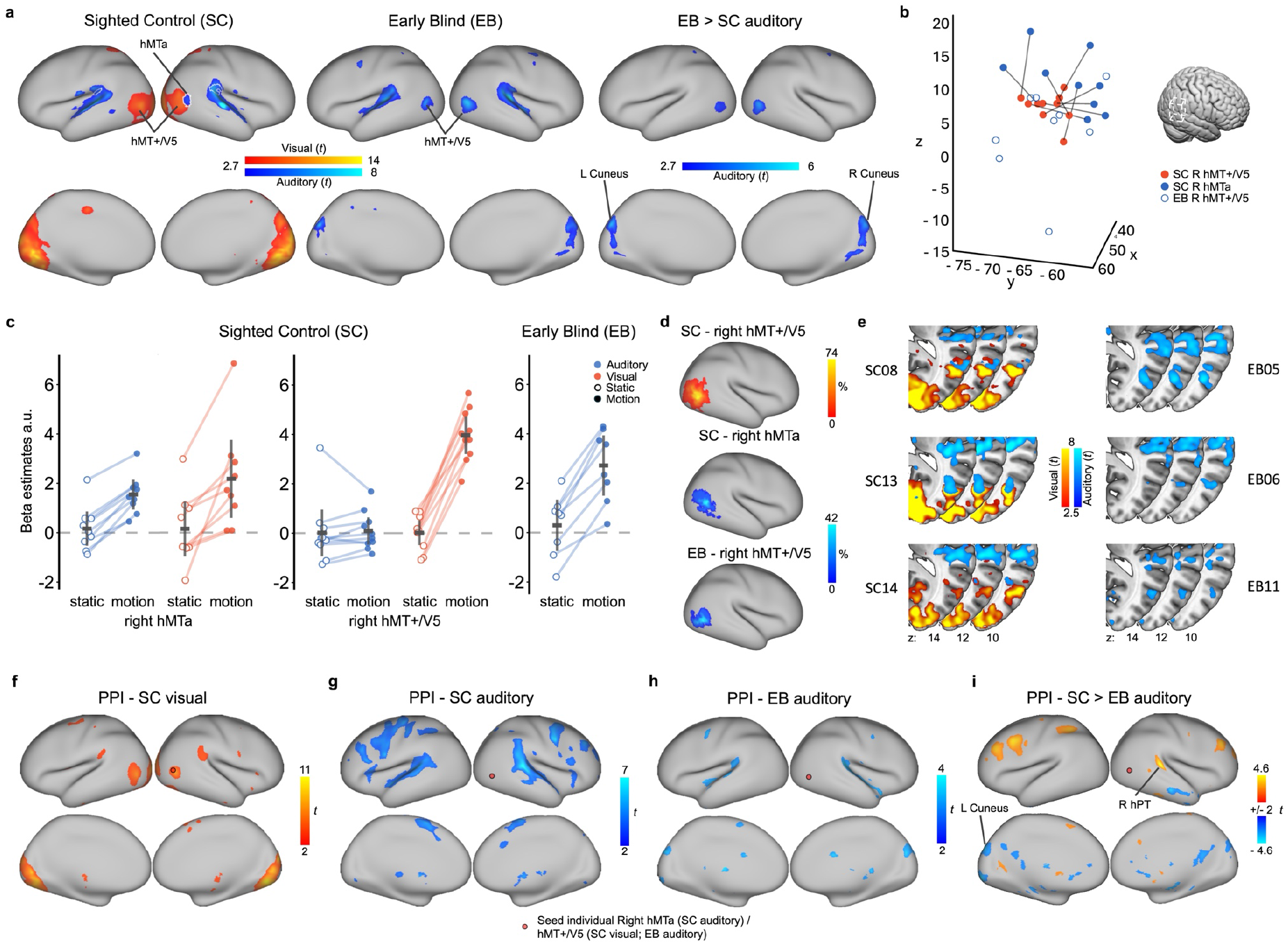
Univariate results investigating the location of motion selectivity in sighted and blind individual at the whole brain scale. a, Group level results (*t*-values) showing motion selectivity (contrast [motion>static]) from the visual (sighted controls) in red and auditory (in sighted and blind individuals) in blue (left and centre figures). Group comparison results (*t*-values) showing auditory motion-enhanced activity in blind individuals from the auditory motion localizer (contrast EB > SC [motion>static]) (right figure). Statistical maps are shown at a voxel-wise p < 0.01 (uncorrected) for visualization purposes. **b,** 3D plot of the peak of activation in MNI coordinates of the individually defined regions of interest. Solid lines connect visual and auditory motion fROIs of sighted controls. **c,** Beta estimates extracted from each individually defined fROI and modality for the static and motion condition reported for illustration purpose only, a.u. = arbitrary units. **d,** Group results showing the percentage of overlap between the individually defined regions of interest (fROI) in each group and modality. **e,** Examples of motion selectivity activation at the single subject level. **f-i,** Group-level Psychophysiological interaction (PPI) analyses revealing motion-specific changes in functional coupling between individually defined right hMTa (SC auditory) and right hMT+/V5 (SC visual; EB auditory) and the rest of the brain based on the functional localizer dataset. The results show connectivity patterns in sighted controls during the auditory motion localizer **(f),** early blind individuals during the auditory motion localizer **(g)** sighted controls during the visual motion localizer **(h),** and direct comparison between sighted and early blind groups highlighting regions showing enhanced motion-dependent coupling with hMT+/V5 during auditory motion processing **(i).** PPI statistical maps are shown at a voxel-wise p < 0.05 (uncorrected) for visualization purposes.

### Visual and auditory motion layer profile of hMT+/V5 in sighted and blind individuals

Next, we characterized the layer profiles of hMT+/V5 of sighted and early blind individuals while attending moving stimuli. Ultra–high-resolution fMRI at 0.75 mm isotropic combined with advanced sequence of acquisition (SS-SI VASO; Huber et al., 2014) requires a restricted field of view, which in the present experiment was limited to the middle temporal region in the right hemisphere. This choice is supported by prior evidence that multisensory responses to both visual and auditory motion are stronger in right hMT+/V5 (Dormal et al., 2016; Rezk et al., 2020). To test the feedforward versus feedback organization in hMT+/V5 for each group and modality, we focused on the individually defined fROIs and the individual layer segmentations (see Methods and Fig.3a). For each fROI, we extracted voxel-wise GLM statistical t-values and plotted the average as a function of cortical depth for BOLD and VASO acquisitions. Although the SS-SI-VASO signal is less sensitive than BOLD, its enhanced laminar specificity revealed distinct laminar response profiles across groups and conditions, as shown by comparing the across layer VASO response structure to models of feedback and feedforward activity profiles (see Methods). To assess the feedback/feedforward nature of these responses quantitatively, we compared each laminar profile to models derived from a quadratic polynomial function (“U-shaped” for feedback, “inverted-U-shaped”, and “skewed inverted-U-shaped” for feedforward). We additionally included an asymmetric, superficial-biased (“skewed inverted-U-shaped”) model to capture the expected laminar profile of feedforward-dominated processing at the spatial resolution of layer-fMRI. Although feedforward afferents primarily terminate in layer IV (Felleman & Van Essen, 1991), activity rapidly propagates through the canonical cortical microcircuit to supragranular layers II/III via ascending intracortical connections, resulting in a response that is not expected to remain confined to the granular layer (Callaway, 1998; Douglas & Martin, 1991). Furthermore, the temporal integration inherent to fMRI likely captures not only the initial feedforward input but also early recurrent interactions within the local cortical circuit and reciprocal exchanges with neighbouring visual areas. Together with residual superficial vascular contributions that may persist even in BOLD-corrected VASO acquisitions (Huber et al., 2014; Lankinen et al., 2022), these considerations motivated the use of an asymmetric feedforward model. Importantly, the asymmetric model was independently supported by the laminar response evoked by visual motion in hMT+/V5 of sighted participants (Fig.3 top-center). Because visual motion is conveyed to hMT+/V5 through the canonical feedforward visual hierarchy, the finding that this condition was best described by the skewed feedforward model provides an empirical validation of the model, rather than relying solely on anatomical or methodological considerations.

**Figure 3.**
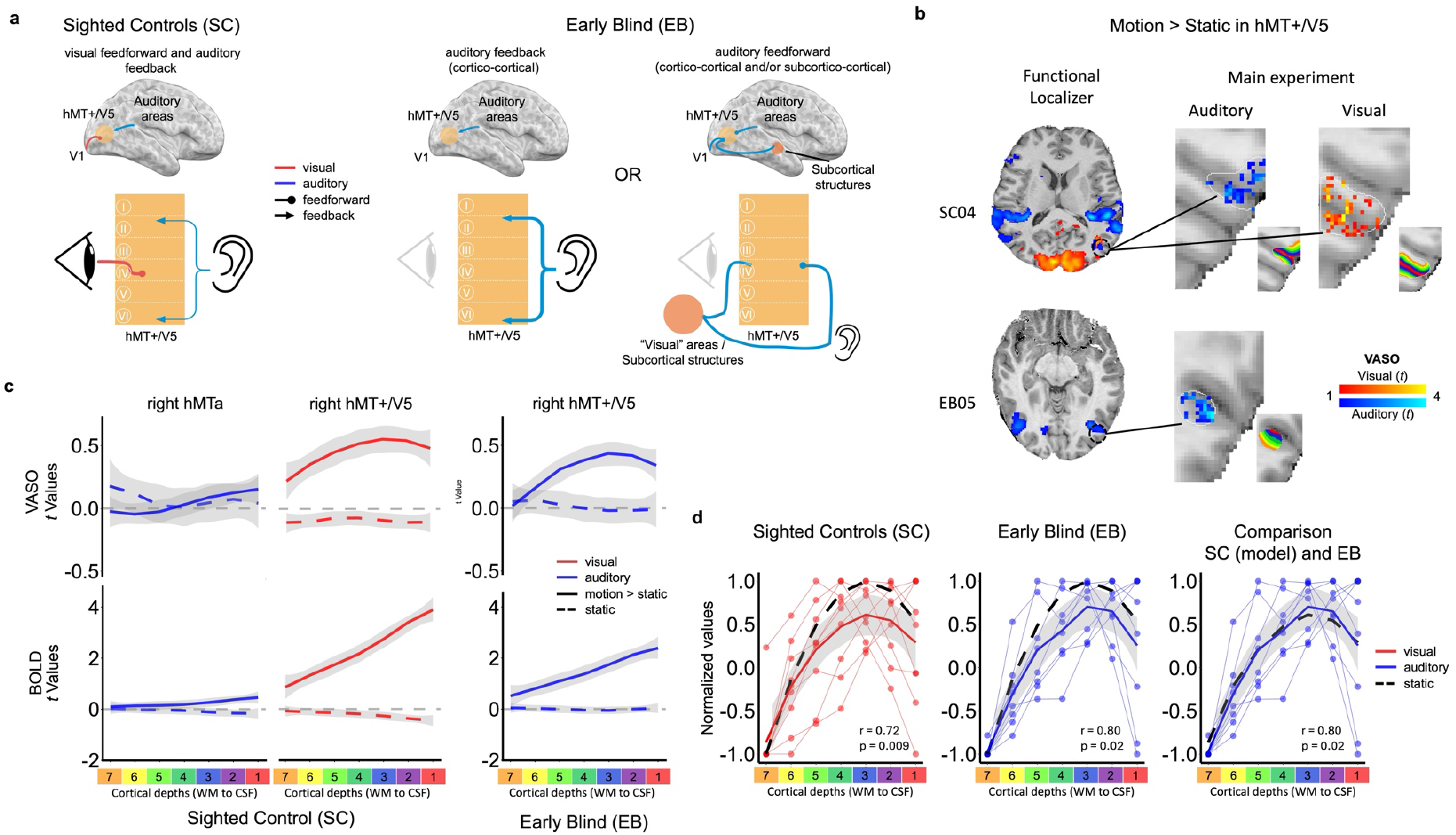
Cortical depth–dependent motion selectivity in sighted and early blind individuals. a, Conceptual models illustrating the expected laminar activation profiles based on putative feedforward and feedback architectures, as a function of sensory modality and visual experience. **b,** Schematic illustration from two representative participants showing the combination of individually defined regions of interest (fROIs), voxel-wise statistical maps from the VASO main experiment, and cortical depth segmentation within each fROI. Seven equidistant cortical layers are shown restricted to the subject-specific fROIs. **c,** Layer-resolved activation profiles within hMT+/V5 and hMTa for the contrast [motion>static] (solid line) and [static] (dashed line) plotted across cortical depth for each group and modality. VASO (top) and BOLD (bottom) results are shown separately. Lines represent the group mean across participants, while shaded regions denote the corresponding 95% confidence intervals, computed using a “loess” smoothing function applied solely for graphical visualization of the confidence intervals to facilitate visualization of laminar trends and was not used for statistical analyses or model fitting. Individual-subject laminar profiles corresponding to each condition are provided in Figs.S4–S6. **d,** Permutation-based model comparison of normalized laminar profiles for the [motion>static] contrast. Observed profiles were compared with a quadratic feedforward template (“skewed inverted-U-shaped”; left, center) and, for early blind participants, with an empirical feedforward model derived from sighted controls (right).

Models and participants laminar profiles were normalized to the range [−1 1] to preserve profile shape while removing inter-individual amplitude differences. Inspection of the normalized profiles revealed that one sighted participant (SC12) and one early blind participant (EB07) exhibited depth-dependent patterns that markedly deviated from the distribution observed within their respective groups and were subsequently evaluated using a standardized deviation criterion (see Methods). A formal assessment confirmed that only these participants had layer values >3 SD relative to the group distribution (SC12: layer 3; EB07: layers 7, 3, 2; Fig.S3). These participants were therefore excluded prior to model-based analyses. We then performed a 100,000-iteration label-randomization test, in which laminar depth labels were permuted within each subject, and the resulting profiles were correlated with the feedforward and feedback models. To ensure adequate sampling across laminar bins and to rule out group differences that could bias depth-dependent analyses, we quantified cortical thickness in all fROIs. Mean thickness across fROIs was ∼2.8 mm (voxel size: 0.75 mm isotropic), allowing at least one voxel per deep, middle, and superficial compartment. Crucially, cortical thickness did not differ between groups (all p > 0.11; see Fig.S12 and Method S3).

In sighted controls, we investigated whether visual and auditory motion elicit feedforward and feedback activation (Fig.3b left). The VASO-derived laminar profile of visually defined hMT+/V5 during visual motion ([motion>static]) increased from deep layers and peaked in mid-cortical depths (Fig.3c top-center), a pattern that was also evident across individual participants (Fig.S4), and showed strong correlations with the feedforward “skewed inverted-U-shaped” model (r = 0.72, p = 0.009; Fig.3d left), a weaker positive correlation with the “inverted-U-shaped”, which did not reach significance (r = 0.46, p = 0.15; Fig.S9 center) and a negative correlation with the feedback “U-shaped” model (r = −0.46, p = 0.85; Fig.S8 center). Importantly, this mid-layers enhancement in hMT+/V5 was specific to motion since extracting separate layer profiles for [motion] and [static] conditions showed that only motion stimuli produced the characteristic mid-layer peak, whereas static stimulation elicited no laminar modulation in VASO (Fig.S5 and Fig.S6). Conversely, the auditory-defined hMTa in sighted individuals exhibited a weaker response to moving sounds that was maximal toward superficial layers during auditory motion stimulation (Fig.3c top-left). Consistently, auditory responses in hMTa of sighted participants exhibited a qualitatively superficial-biased depth profile, as expected for feedback-dominated processing. However, owing to the comparatively small response amplitude, correlations with the predefined feedback model (“U-shaped”) did not reach statistical significance (r = 0.03, p = 0.45; Fig.S8 left), neither with the feedforward models (“inverted-U-shaped”: r = −0.03, p = 0.54; Fig.S9; “skewed inverted-U-shaped”: r = 0.13, p = 0.43; Fig.S10).

BOLD responses to auditory and visual [motion>static] in sighted controls showed an expected monotonic increase toward superficial layers (Fig.3c bottom −left and −center; see Fig.S4 for single subject layer profile), the characteristic supragranular bias attributed to ascending veins in BOLD. As expected, neither auditory nor visual static stimuli elicited measurable BOLD responses across cortical depth (Fig. 3c bottom-left and −center; Fig. S6). The absence of depth-dependent activity under static conditions confirms the motion selectivity of the functional ROIs and provides an important internal control, demonstrating that the laminar analysis does not generate spurious depth-dependent responses in the absence of task-related activation. Then, to assess whether laminar profiles reflected activity reliably greater than baseline, we performed non-parametric Wilcoxon signed-rank tests (one-sided, μ = 0) on specific layer VASO responses. To reduce the burden of multiple comparisons while preserving sensitivity, layers were down-sampled into three depth bins (deep, middle, superficial; see Methods). After false-discovery-rate correction across depth bins, visual motion responses in hMT+/V5 of SC were significantly greater than zero in each depth bin (deep, middle and superficial: V = 55, adjusted-p < 0.001; Fig.S11 center). Auditory motion responses in hMTa displayed the expected superficial predominance, with positive responses confined to superficial layers (V = 31, uncorrected-p = 0.039; Fig.S11 left), although this effect did not survive correction for multiple comparisons across layers. For completeness and transparency, both corrected and uncorrected statistics are reported (Table S3).

We next asked whether auditory motion reaches hMT+/V5 in early blind individuals via a feedback or a feedforward circuitry (Fig.3a center and left). The VASO laminar profile in EB participants displayed a clear feedforward-like pattern (Fig.3c top-right; consistently observed across individual participants see Fig.S4), additionally confirmed by a strong correlation with the feedforward “skewed inverted-U-shaped” model (r = 0.80, p = 0.02; Fig.3d center), and a weaker positive correlation with the “inverted-U-shaped” (r = 0.48, p = 0.13 Fig.S9 right) and negative with the feedback “U-shaped” model (r = −0.48, p = 0.87; Fig.S8 right). This laminar profile closely matches the mid-layer peak observed for visual motion in sighted individuals. To further test whether the EB laminar profile matched that of visual motion in sighted individuals, we used the mean SC profile as a model in an additional permutation analysis across EB participants. This analysis confirmed that EB responses closely matched the sighted feedforward profile (r = 0.80, p = 0.02; Fig.3d right).

Similar to the sighted in vision, the BOLD layer profile to moving sounds in EB increased from white-matter boundaries toward superficial layers (Fig.3c bottom-right). The feedforward modulation was indeed specific to motion only, as static sounds produced no response, neither in VASO nor in BOLD (Fig.3c right; Fig.S6) again confirming the motion selectivity of our fROIs. One-sample Wilcoxon signed-rank tests confirmed that auditory motion responses in EB were also significantly greater than zero in each depth bin (deep: V = 40, adjusted-p = 0.019, middle and superficial: V = 45, adjusted-p = 0.003), demonstrating that this profile reflects reliable task-evoked activity (Fig.S11 right, Table S3). Together, these analyses showed that crossmodal auditory responses to motion in the blind hMT+/V5 display a feedforward laminar profile comparable with visual motion processing in sighted controls.

### Psycho-Physiological-Interaction analyses (PPI) of hMT+/V5 in sighted and blind individuals

While laminar profiles reveal whether motion-related signals reach hMT+/V5 through feedforward or feedback circuitry, they do not identify where these inputs originate, especially given the restricted field of view required by our laminar recordings. To determine the large-scale networks supplying task-dependent input to right hMT+/V5, we performed whole-brain connectivity analyses using Psycho-Physiological Interactions (PPI) based on the localizer dataset, which provided full brain coverage, unlike the high-resolution laminar acquisition. PPI quantifies how task-related fluctuations in a seed region covary with activity in other brain regions. For each participant, we extracted the first eigenvariate of the hMT+/V5 fROI (psychophysiological regressor), capturing the oscillation between [static] and [motion] conditions, and tested motion-selective voxel-wise connectivity changes across the whole brain. Analyses were conducted separately for each group-and-modality-specific fROI (SC: visually defined right hMT+/V5 and auditory-defined right hMTa; EB: auditory-defined right hMT+/V5).

In sighted controls, visually defined right hMT+/V5 showed increased task-dependent coupling with right lingual gyrus and right calcarine sulcus (V1) during visual motion processing (Fig.2f; Table S5). Comparing auditory-motion connectivity across groups revealed a clear dissociation: sighted individuals showed strengthened coupling with the auditory cortex and in particular, the Planum Temporale known to engage in auditory motion processing (Battal et al., 2019), whereas early blind individuals instead exhibited stronger connectivity with the left cuneus (Fig.2i; Table S5). This pattern aligns with the laminar results, supporting the interpretation that auditory motion reaches hMT+/V5 in blind individuals via a feedforward pathway hypothetically originating from the early visual cortex.

## DISCUSSION

Revealing the mesoscale organization of crossmodal responses in the occipital cortex of congenitally blind individuals is critical for understanding how sensory deprivation shapes the development of sensory circuits. We characterized layer-specific responses to moving and static visual and auditory stimuli in individually and functionally defined hMT+/V5 in sighted and early blind individuals.

We first show that visual motion responses in sighted hMT+/V5 are best captured by a feedforward architecture, providing an empirical benchmark for interpreting auditory responses after blindness. Against this benchmark, our study reveals that hMT+/V5 in early blind individuals exhibits a remarkably similar circuit-level response to auditory motion, effectively maintaining a canonical feedforward laminar organization despite crossmodal plasticity. Importantly, the observed laminar effects were specific to moving stimuli, as auditory and visual static stimulation failed to elicit comparable depth-dependent responses, demonstrating that the identified profiles reflect sensory motion processing rather than non-specific laminar biases.

This feedforward laminar pattern of functionally selective response to moving sounds suggests that crossmodal plasticity in early blindness does not chiefly reflect a scaling-up of top-down feedback signals from higher-order association cortex (Macaluso & Driver, 2005; Merabet & Pascual-Leone, 2010) as previously suggested (Büchel, 2003; Klinge et al., 2010; Rons, 1996). Instead, our results show that auditory inputs engage hMT+/V5 mainly through reorganization of feedforward pathways, possibly involving strengthened inputs from early visual (e.g. V1) or subcortical structures (e.g. LGN). These results are supported by previous electrophysiological studies in humans showing extremely rapid response (as early as 35ms post-stimulus) to crossmodal inputs in the occipital cortex of EB (Leclerc et al., 2000; Müller et al., 2019) or by transcranial magnetic stimulation studies showing altered crossmodal processing after V1 interference early post-stimulus presentation (Matuszewski et al., 2025).

The finding that moving sounds evoke a feedforward-like laminar response profile in hMT+/V5 in early blind individuals suggests that auditory motion information engages this region through developmentally reorganized ascending pathways. In sighted people, hMT+/V5 is embedded within a predominantly feedforward motion-processing hierarchy and receives the majority of its driving input from early visual cortex, particularly V1 (Felleman & Van Essen, 1991). Anatomical tracing studies in non-human primates demonstrate that V1 constitutes the largest cortical source of afferent input to area MT, accounting for approximately 60–70% of its total cortical inputs (Desimone & Ungerleider, 1986; Felleman & Van Essen, 1991; Maunsell & Van Essen, 1983; Rockland, 1989, 1995). Consistent with these anatomical findings, electrophysiological recordings show that visual responses in MT are strongly dependent on intact V1 input: lesions of V1 abolish or severely attenuate direction-selective responses in MT for the majority of neurons, with residual activity confined to a minority of cells driven by alternative subcortical pathways (Movshon & Newsome, 1996; Ponce et al., 2008). Furthermore, human neuroimaging and lesion studies converge on this hierarchical organization by showing strong structural and functional coupling between V1 and hMT+/V5, and that damage to early visual cortex markedly reduces visually evoked responses in hMT+/V5, despite the preservation of higher-order visual areas (Ajina et al., 2015; Bridge et al., 2008). Together, these findings establish that, under typical developmental conditions, hMT+/V5 operates largely as a downstream receiver of feedforward signals mainly originating in early visual cortex. One plausible contributor of middle layers activity by moving sounds in early blind people is therefore the early visual cortex, which is known to exhibit robust auditory responsiveness following early visual deprivation (Collignon et al., 2011). Consistent with this interpretation, our PPI analyses reveal task-dependent interactions between hMT+/V5 and early visual regions in the blind group, indicating that, despite lack of visual inputs during development, crossmodal auditory processing can recruit feedforward circuitry typically associated with visual motion processing. Evidence from animal models demonstrates that early blindness induces profound reorganization at subcortical stages of sensory processing (Bronchti et al., 1989; Chabot et al., 2008; Kaiserman-Abramof et al., 1980; Karlen et al., 2006). Auditory responses have been observed in visual thalamic nuclei, including the dorsal lateral geniculate nucleus (dLGN) following congenital blindness or enucleation, providing a potential route for auditory signals to reach the early visual cortex via ascending pathways (Bronchti et al., 1989; Charbonneau et al., 2012). Such subcortical changes are accompanied by large-scale alterations in thalamocortical and corticocortical connectivity, as documented in animal studies after early visual deprivation (Dehay et al., 1996; Karlen et al., 2006; López-Bendito et al., 2022). Although reduced in size, the LGN and the optic radiations relaying information to V1 are anatomically present in blind humans (Bridge et al., 2009; Cecchetti et al., 2016; Ptito et al., 2008). Since the LGN does not receive retinal inputs in EB, these connections may support the transfer of auditory information through alternate connections from auditory thalamus (MGN) to area V1 via the LGN. This is supported by animal studies showing LGN activation upon auditory stimulation in the blind mole rat (Bronchti et al., 2002; Doron & Wollberg, 1994). Together, these findings raise the possibility that the feedforward-like auditory responses we observe in hMT+/V5 originate, at least in part, from a reconfiguration of ascending sensory pathways to primary visual areas from subcortical areas such as LGN, MGN, and the Pulvinar (Ewall et al., 2021). An alternative route, not mutually exclusive with the one just described, would be that moving sounds reach the middle temporal cortex of EB using a direct connection between the LGN and pulvinar with hMT+/V5, a pathways that exists in non-human animals (Berman & Wurtz, 2011; Born & Bradley, 2005) and sighted humans (Ajina & Bridge, 2018; Allen et al., 2015; Arcaro et al., 2015; Gaglianese et al., 2012; Leh et al., 2008).

In contrast to EB, laminar auditory-motion responses in sighted individuals preferentially engaged superficial layers and connectivity analyses revealed enhanced auditory motion selective coupling with motion-selective auditory regions including the Planum Temporale, putatively involving a feedback-dominated modulation (Macaluso et al., 2000; Macaluso & Driver, 2005). Superficial layers receive top-down contextual information and have been shown to exhibit modulation of activity by other sensory modalities in sighted non-human animals (Chou et al., 2020; Ibrahim et al., 2016; Iurilli et al., 2012, p. 202; Lakatos et al., 2007). Activity in superficial layers in hMTa for moving sounds in sighted individual was weak compared to the activity observed in blind people for moving sounds or in sighted people for moving visual stimuli (see Fig. S11). This reduced statistical sensitivity is expected given the substantially smaller response amplitudes associated with feedback-driven activity compared with feedforward sensory input (Lawrence et al., 2019; Pizzuti et al., 2025) and reflects the low amplitude response also expressed in the whole brain results from the auditory motion localizer in sighted controls (Fig.3 c left).

Interestingly, across primary sensory cortices, thalamocortical synapses to layer 4 (L4) have an early critical period for plasticity (Barkat et al., 2011; Crair & Malenka, 1995; Feldman et al., 1998; B. Jiang et al., 2007), but synapses from L4 to L2/3 undergo plasticity through adulthood (B. Jiang et al., 2007). This leads to the intriguing possibility that crossmodal recruitment by sounds in the visual cortex in late acquired blindness (Mattioni et al., 2022), after the development of the visual system, may be expressed by reorganizing feedback laminar circuitry in contrast to early acquired blindness (Collignon et al., 2013).

Together, these findings provide evidence for a feedforward crossmodal remapping of motion processing in the absence of vision. They highlight a remarkable capacity of cortical circuits to incorporate non-visual inputs while preserving canonical microcircuit organization, extending previous demonstrations of macroscale crossmodal activation to a mesoscale, laminar level (Battal et al., 2019; Collignon et al., 2011; Mattioni et al., 2022; Merabet & Pascual-Leone, 2010). Importantly, our results reconcile two perspectives: while multisensory input in the occipital cortex in sighted individuals seems feedback-driven (Iurilli et al., 2012; Macaluso et al., 2000; Macaluso & Driver, 2005), early sensory deprivation enables low-level auditory inputs to drive hMT+/V5 in a feedforward manner, effectively converting the visual motion complex into a modality-flexible motion processor (Amedi et al., 2017; Ricciardi et al., 2007). It would be interesting to see if such reorganized feedforward pathways that convey sounds to hMT+/V5 link to the observed reduced coding of auditory motion in the Planum Temporale of early blind people (Battal et al., 2019; F. Jiang et al., 2016).

These findings suggest that canonical cortical circuits maintain hierarchical input specificity even under conditions of sensory deprivation (here motion selectivity), and that crossmodal plasticity can reorganize input pathways while preserving computational principles. Additionally, these findings have implication for sight restoration developments as reorganization of bottom-up sensory transduction processes following early blindness might impact the transfer of the regained visual input (Collignon et al., 2012, 2015; Mattioni et al., 2025). The profile of this reorganization is expected to be different in late blind people (Mattioni et al., 2022). Finally, our work demonstrates the unique opportunity of layer-resolved fMRI to reveal experience-dependent plasticity of the brain circuitry, providing a mechanistic bridge between human non-invasive imaging and invasive animal studies of laminar computation (Huber et al., 2017, 2019). With advances in fMRI technology and the increasing availability of ultrahigh-field human MRI, layer-specific fMRI will play an expanding role in complementing whole-brain high-resolution fMRI to provide deeper insights into neural circuit changes due to sensory deprivation (Chai et al., 2024; Koiso et al., 2022).

## METHODS

### Subjects

Nine early blind adults (2 women; mean age = 35 years, SD = 14.1) and 10 sighted control participants (4 women; mean age = 29.5 years, SD = 6.19) took part in the experiment. All participants were right-handed except for one sighted and three blind individuals, who identified as ambidextrous. The groups did not significantly differ in terms of sex distribution (Fisher’s exact test: p = 0.63) or age (two-sample t-test: *t*(16) = 0.44, p = 0.66). However, there was a significant difference in educational background, with sighted participants having spent more years in formal education on average (early blind: mean = 13.9, SD = 1.9; sighted controls: 16.8, SD = 2.39; two-sample t-test: *t*(16) = −2.95, p = 0.009). Additional participant details are provided in Table S1. All individuals reported normal hearing and no history of neurological disorders or brain injury. Blindness in the blind group was due to peripheral causes, and sighted controls had normal or corrected-to-normal vision. Informed consent and MRI safety assessments were completed before participation, and all the participants received financial compensation. The study was approved by the Research Ethics Committee of Cliniques Universitaires Saint-Luc of the Université Catholique de Louvain (UCLouvain), and Hospitalo-Facultaire of the University of Liège. Most participants completed three MRI sessions as part of a larger data collection involving other experiments.

### MRI acquisition

Data acquisition was performed on a whole-body MAGNETOM Terra 7T (Siemens Healthinneers, Erlangen, Germany) at the GIGA-Cyclotron-In Vivo Imaging Research Centre at the University of Liège (Belgium) using a 32-channel RX head-coil (Nova Medical, Wilmington, MA, USA).

#### Anatomical image

Anatomical images have been collected in each session (3 sessions for most of the participants) using an MP2RAGE sequence (Marques et al., 2010). During each session, we acquired two whole-brain anatomical images with different inversion times: TR = 4300 ms, TE = 2.27 ms, echo spacing = 7.2 ms, excitation TR = 4.54 ms, 192 slices, 0.75-mm isotropic voxels, matrix size = 192 × 300 × 320, flip angles = 4°/4°, GRAPPA factor = 3, and inversion times TI₁ = 1000 ms and TI₂ = 3200 ms.

#### Visual and auditory motion localizer

Whole brain functional images were acquired using a 1.6 mm isotropic resolution with a BOLD-GE acquisition: number of slices = 72, TR = 1.8 s, TE = 22 ms, FA = 70°, MB = 2, FoV = 214.4 x 217.6 x 115.2 mm, number of volumes = 264 and 414 (visual and auditory localizers respectively). The first 4 volumes of each scan were discarded to allow the MR signal to reach steady-state magnetization.

#### Main Experiment

For the main experiment, we acquired CBV-sensitive data using a 7T-optimized Vascular Space Occupancy (VASO) protocol. We implemented the slice-selective slab-inversion (SS-SI VASO) scheme combined with a 3D EPI readout (Huber, Finn, et al., 2021; Stirnberg & Stöcker, 2021), enabling nominal 0.75 mm isotropic resolution suitable for sub-millimetre analyses. The acquisition covered a 130.5 × 130.5 x 18 mm sagittal field of view (174 × 174 matrix) across 24 slices. Key parameters included TE = 24.7 ms, volume TR of 1.711 s, time of acquiring pairs of BOLD and VASO TR of 3.8s, a variable-flip-angle excitation (FA = 32.8°-60°) to reduce T1-related blurring, and 6/8 partial Fourier). The inversion time was set to 1515.6 ms to null intravascular signal, and acceleration was achieved with dual-polarity GRE-GRAPPA (factor 3) reference lines. The sequence was developed in the IDEA environment (VE12U-SP01) and is distributed as a C2P package via the Siemens Teamplay platform. The first 4 volumes of each scan were discarded to allow the MR signal to reach steady-state magnetization.

### Task paradigm

#### Visual stimuli

Visual stimulation was delivered via a projector (Sanyo, PLC XT21/L) and the image was reflected by two mirrors before reaching the participants’ eyes (one mounted over the MRI coil and one in between the projector and the coil mirror). In all the visual experiments, Random Kinematic Dots (RDK) were presented covering the whole field of view. In each frame under any conditions, white dots with 0.1° diameter were presented on a black background. In both the visual localizer and the visual main experiment, we presented alternating moving or static dots with identical characteristics. Moving dots had a speed of 7°/s moving coherently in one of four directions per trial (up-, right-, down-, or left-ward) and a screen lifetime of 200 ms with starting position randomly assigned. Static dots had a lifetime equal to the entire static block duration of the specific experiment, and a random location was assigned at the beginning of each static block.

#### Auditory stimuli

To generate ecological auditory motion percepts and ensure precise spatial localization inside the MRI environment, we first recorded all auditory stimuli individually for each participant using binaural in-ear microphones. Because each person’s pinnae are unique, and because the filtering they impose on sound directionality is learned by each person from early childhood, the use of individual recordings is necessary to avoid perceptual confusion (Møller et al., 1996). These recordings were acquired in a semi-anechoic chamber during a dedicated pre-scan session. Participants were seated at the center of a custom sound-delivery setup, with their head stabilized on a chin rest and oriented toward two semicircular arrays, one horizontal and one vertical, each composed of 31 loudspeakers. Both arcs had a radius of 1.1 m, ensuring a constant 1.1 m distance between every loudspeaker and the participant’s ears (Fig.1a left). The horizontal array was aligned with ear level, while the vertical array was positioned along the mid-sagittal plane.

To generate smooth, continuous auditory motion, pink noise was partitioned into 31 equal temporal segments and sequentially presented through corresponding loudspeakers, with no temporal gap or overlap between segments. Stimulus duration was 0.85 s (2.71 m/s) for the auditory localizer (0.425 s, 5.42 m/s in the behavioral task) and 0.80 s (2.88 m/s) for the main experiment, with 50 ms rise/fall times applied to each stimulus. This procedure produced a seamless percept of motion at a constant 65 dB from the listener’s position. We generated four translational motion trajectories, upward, downward, rightward, and leftward, all spanning 120° of participants’ peripheral space.

A static condition was also recorded using a mannequin (3Dio, Free Space Pro), delivered from the central loudspeaker located at the intersection of the two orthogonal arcs. Static sounds matched the duration of motion events and included pink noise (standard trials) and white noise (target trials).

All moving sound recordings were captured using a Zoom H4n recorder −200 m in combination with Master Series Sound Professionals TFB-2 in-ear microphones and were subsequently replayed in the MRI scanner during both the auditory motion localizer and the main experiment. By convolving the stimuli with each participant’s own pinna- and head-related transfer functions, this individualized recording procedure produced an immersive, externalized representation of auditory space, closely approximating real-world auditory motion.

#### Visual motion localizer

Participants completed a ∼5-minute run during which they were presented with blocks of moving and static dots. Moving dot blocks lasted 10.3 seconds and contained white dots moving along the horizontal axis; each of the six trials presented leftward and rightward motion consecutively. Static dot blocks lasted 9 seconds. Blocks were separated by an inter-block interval (IBI) of 5.4 seconds, and a total of 15 cycles were presented. Participants were instructed to fixate a central red cross and perform two tasks: (1) press a button whenever the fixation cross changed color, and (2) indicate when the direction of motion repeated across consecutive moving-dot trials or when the static dots changed color from white to gray within a trial. See Fig. S2 and Method S1 for more information regarding the behavioral performance during the task.

#### Auditory motion localizer

Analogous to the visual motion localizer, participants completed a ∼10-minute run during which they were presented with blocks of moving and static sounds along the horizontal axis. Each of the six trials contained leftward and rightward-moving sounds consecutively, and a total of 22 cycles were presented. Participants were instructed to indicate trials in which a static sound was a different one (white noise instead of the regular pink noise), and in auditory motion blocks when the trial was perceived to be moving faster within a brief interval. Participants were instructed to keep their eyes closed during the whole run. See Fig. S2 and Method S1 for more information regarding the behavioral performance during the task.

#### Main Experiment

The main experiment included a visual version for sighted participants and an auditory version for both sighted and blind participants, using an identical design across modalities. Participants completed four runs per modality: sighted participants completed eight runs in total, alternating modalities and counterbalancing the modality of the first run across participants, while blind participants completed four auditory runs. Each run lasted 9 minutes. Each run comprised alternating blocks of static and moving stimuli, with 8 static and 8 moving blocks per run. Each block contained 12 trials of 1.6 s, and the block sequence was fixed across runs, beginning with a static block to facilitate averaging across runs and enhance the signal for analysis.

In the visual modality, static blocks displayed stationary dots, whereas moving blocks presented dots moving along horizontal or vertical axes (e.g., upward and downward within a trial) and the presentation of a specific axis of motion equally distributed along the run. In the auditory modality, static blocks presented continuous sounds within a trial, while moving blocks contained sounds perceived as moving along horizontal or vertical axes.

Participants performed an attention task: during visual runs, they responded when the fixation cross changed color from red to green, when dots changed color during static blocks, or when a motion direction was repeated within a moving trial (e.g., two consecutive upward movements). During auditory runs, participants responded to changes in static sounds (from pink to white noise) or repeated motion directions within moving trials. Participants were instructed to keep their eyes closed throughout auditory runs. See Fig. S3 and Method S2 for more information regarding the behavioral performance during the task.

The stimulation of all the experiments were delivered in Matlab (Mathworks, Inc.), using the Psychophysics Toolbox extensions (Brainard, 1997; Kleiner et al., 2007; Pelli, 1997).

### Structural and functional image processing and statistical analyses

See Fig.S1 for an overview of the analyses pipeline.

#### Localizers for regions of interest (fROI) definition

Structural and functional whole brain data at 1.6 mm isotropic resolution were pre-processed and statistically analysed using a standard approach implemented in bidspm (v3.0.0; https://github.com/cpp-lln-lab/bidspm; Gau et al., 2022) (which is wrapper application for the software Statistical Parametric Mapping (SPM12 - v7771; Wellcome Center for Neuroimaging, London, UK; https://www.fil.ion.ucl.ac.uk/spm; RRID:SCR_007037) to ease fMRI analyses of Brian Imaging Data Structure (bids) format data (Gorgolewski et al., 2016). Analyses have been performed on Matlab 2014a (Mathworks, Inc.) installed on a Linux-Ubuntu (Ubuntu 22.04.5 LTS, Jammy Jellyfish) computer. Preprocessing was performed both in individual and standard (MNI) space in order to, in the first case, extract fROIs to be used to characterize their layer profile, where it is more suitable to work in individual space to not alter the data space, and the second one, perform group-level analyses to evaluate motion selectivity and connectivity at the whole brain scale. We followed these steps: slice-timing correction was applied using the middle slice (36th) as reference (sinc interpolation), followed by realignment using the mean functional image (6 degrees of freedom; least-squares cost function). The anatomical image was bias-field corrected, segmented, and normalized to MNI space (IXI549 template; 1-mm resolution; 4th-degree B-spline interpolation) using unified segmentation. Tissue probability maps were used to skull-strip the anatomical image by removing voxels with p(GM)+p(WM)+p(CSF) < 0.75. The mean functional image was co-registered to the bias-corrected anatomical scan (6 degrees of freedom; normalized mutual information), and the resulting transformation was applied to all functional volumes. Deformation fields from segmentation were then applied to normalize all functional images to MNI space (native functional resolution; 4th-degree B-spline interpolation). Finally, normalized and not normalized functional data were spatially smoothed using a 3D Gaussian kernel (FWHM = 4 mm for fROI definition in individual space; 6 mm for group level results in MNI space). Statistical analyses were performed as follow: for the visual and auditory motion localizers, we estimated a voxel-wise general linear model (GLM) that included separate regressors for static and motion stimulation. Each regressor was convolved with the canonical hemodynamic response function implemented in SPM. To account for residual movement, the 6 rigid-body motion parameters (three translations and three rotations) were added as nuisance regressors. For each participant and statistical results in individual space with 4 mm kernel smoothing, we computed the [motion>static] contrast to delineate regions preferentially engaged by motion in the corresponding modality. The resulting fROI masks were visually inspected and subsequently transformed to the functional VASO. Specifically, fROIs were morphologically distorted to the EPI space using antsApplyTransforms (ANTs, v2.6.2; Tustison et al., 2021), with the transformation matrix derived from the alignment between the high-resolution structural image (UNIT1) and the T1-weighted image derived from the VASO functional data. This ensured that fROI boundaries were accurately mapped to the native functional space used for depth-resolved analyses. The procedure used to derive and align the UNIT1 anatomical volume to the VASO-based T1-weighted image is described in detail in a subsequent section. To perform group-level results, we considered the data in MNI space and smoothed at 6 mm kernel, these individual contrast images for [motion>static] were subsequently smoothed with a 6-mm FWHM Gaussian kernel and submitted to a second-level random-effects analysis. Group-level inference relied either on whole-brain family-wise error (FWE) correction at p < 0.05 on the whole brain volume or using small volume correction centred on the coordinate of interest (see Table S2 for details). See Fig.S1a row for an overview summary.

#### PsychoPhysiological Interaction (PPI) analyses

To investigate task-dependent changes in functional coupling associated with motion processing, psychophysiological interaction (PPI) analyses were performed using the general linear model framework in SPM12 (SPM12 - v7771; Wellcome Center for Neuroimaging, London, UK; https://www.fil.ion.ucl.ac.uk/spm; RRID:SCR_007037). PPI analyses were conducted separately for the visual and auditory motion localizers, providing whole-brain coverage.

For each subject, the physiological regressor was defined as the first eigenvariate of the BOLD time series extracted from a seed region of interest (fROI), adjusted for effects of interest. Seed fROIs were defined individually based on subject-specific activation peaks within hMT+/V5 or hMTa, identified from the corresponding localizer contrast [motion>static].

The psychological regressor encoded the localizer contrast of interest (visual or auditory [motion>static]) and was convolved with the canonical hemodynamic response function. The PPI regressor was computed following standard SPM procedures: the BOLD signal from the seed fROI was deconvolved to estimate the neuronal signal, multiplied element-wise with the psychological regressor, and then reconvolved with the canonical HRF. All three regressors (physiological, psychological, and PPI) were included in a first-level GLM along with nuisance regressors, including six motion parameters. Functional images were spatially smoothed using a 6 mm full-width at half-maximum (FWHM) Gaussian kernel prior to model estimation. Subject-level contrast images for the PPI term were entered into second-level random-effects analyses to assess group-specific and group-difference effects. One-sample t-tests identified regions showing significant task-dependent coupling with the seed fROI within each group (sighted controls or early blind), and two-sample t-tests tested for differences between groups. Statistical inference was performed using family-wise error (FWE) correction at p < 0.05 at whole brain and using a small volume correction using a sphere of 10 mm (see Table S5 for more details).

#### Structural preprocessing and cortical depth segmentation

High-resolution anatomical images (UNIT1) derived from the MP2RAGE acquisition were obtained across multiple sessions, rigidly aligned to the first session, and averaged to improve signal-to-noise ratio. These preprocessed data are shared with a companion study currently in preparation and was applied identically across datasets. Non-brain background signal inherent to the MP2RAGE sequence was removed using Presurfer (v1.0; https://github.com/srikash/presurfer). Cortical depth segmentation was performed using layer-MAP (v0.2.0; https://github.com/marcobarilari/layer-MAP; Barilari et al., 2026), which provides an automated framework integrating tools from multiple MRI software packages. The processing pipeline implemented in the toolbox is described below. Anatomical volumes were first corrected for intensity inhomogeneities using the SPM12 bias-field correction algorithm (v7771), as implemented within Presurfer. Tissue classification was then performed using FreeSurfer’s standard surface reconstruction workflow (recon-all, v7.3.0; Fischl, 2012). To enhance the spatial fidelity of the resulting gray- and white-matter boundaries, cortical surfaces were reconstructed in surface space using SUMA (MapIcosahedron, AFNI v24.0.16; Cox, 1996) with a high vertex density (linDepth = 2000). These surfaces were subsequently converted back into volumetric space on an up-sampled grid (3× isotropic up-sampling; 0.25 mm voxel size). To minimize the boundary overlap across adjacent gyri during surface-to-volume projection, a single iteration of morphological growth (dilation followed by erosion) was applied. To enable accurate alignment between anatomically derived segmentations and functional data, the averaged UNIT1 image was registered to a T1-weighted image derived from the VASO functional acquisition (submillimetre distorted EPI space) using ANTs (antsRegistration, v2.6.2; Tustison et al., 2021). The resulting rim masks were visually inspected within the regions of interest to verify segmentation accuracy and subsequently transformed into distorted VASO functional space using antsApplyTransforms (ANTs v2.6.2; Tustison et al., 2021), applying the transformation matrix obtained from the alignment between the structural UNIT1 image and the T1-weighted image derived from the VASO functional data, following the same procedure used to align the fROIs to the VASO functional images. Finally, cortical depth estimates were generated using LayNii (v2.6.0; Huber, Poser, et al., 2021) by subdividing the cortical ribbon into 7 and 3 cortical depths using the “equi-count” method, ensuring an approximately equal number of voxels per depth bin. These layer masks were intersected with individually defined regions of interest in right hMT+/V5 to extract depth-resolved parameter estimates from the BOLD and VASO GLM analyses. See Fig.S1c row for an overview summary.

#### Sub-millimetre SS-SI VASO and BOLD fMRI for layer profile

Functional high-resolution fMRI data with a restricted field of view acquired using SS-SI VASO acquisition require a careful and custom project-based pipeline using several software packages The first step consisted of denoising each scan using NORDIC (Vizioli et al., 2021) optimized for VASO to increase the signal-to-noise ratio. Then, each scan was separated into the odd and even volumes (corresponding into cerebral-blood-volume based (CBV) and BOLD images, corresponding to images with and without blood nulling) and performed motion correction separately in AFNI (3dallineate, AFNI v24.0.16; Cox, 1996) applying a brain mask and using the average of the first 3 volumes of each series as a reference. Final re-aligned images and motion parameters were visually inspected and compared across the two series of blood-nulled and BOLD to confirm that they were similar, participant motion was not too high, and the volume-to-volume displacement was corrected. At this point, the 4 runs per modality were analysed as averaged to further improve the signal-to-noise ratio and as single runs. Each series of data (averaged or not) was then restored to each previous length by interpolating in-between volumes (3dupsample, AFNI v24.0.16; Cox, 1996). After temporal up-sample, a dynamic division of the blood-nulled and BOLD volumes was carried out (LN_BOCO, LayNii v2.6.0; Huber, Benedikt, et al., 2021; Huber et al., 2024) to obtain VASO images with minimized residual BOLD contamination. During the preprocessing we also obtained a T1-weighted image from the functional time series of each participant by computing the inverse coefficient of variation across time. This operation emphasizes the temporally stable component of the VASO signal and attenuates fluctuations related to noise or residual BOLD effects, resulting in a high-contrast reference image suitable for spatial alignment. The resulting image served as the participant-specific anatomical reference during co-registration steps in the preprocessing pipeline. A voxel-wise GLM was then fit on both VASO and BOLD (averaged and not) time series using SPM12 (v7771; Wellcome Center for Neuroimaging, London, UK; https://www.fil.ion.ucl.ac.uk/spm; RRID:SCR_007037), and similar parameters implemented for the previously described steps on the whole brain fMRI data. Finally, since VASO values are inversely related to magnitude of brain response, the resulting t-values for VASO were multiplied by −1 to unify the direction of effects. See Fig.S1b for an overview summary.

#### Laminar profile modelling and permutation-based inference

To assess whether laminar response profiles were consistent with feedforward or feedback architectures, we compared subject-specific depth profiles against predefined model templates using permutation-based statistics. All analyses were conducted in R (version 4.4.2; R Core Team, 2024) using custom scripts and standard statistical libraries. Analyses were performed separately for group, sensory modality, and region of interest. Three canonical quadratic models were defined across seven equidistant cortical depths: a symmetric feedback profile (“U-shaped”), a symmetric feedforward profile (“inverted-U-shaped”), and an asymmetric superficial-biased feedforward profile biased toward superficial layers (“skewed inverted-U-shaped”). The inclusion of the skewed model was motivated by both methodological and biological considerations. Although BOLD-corrected VASO substantially reduces sensitivity to draining-vein effects, residual superficial weighting due to vascular architecture and hemodynamic properties may persist (Huber et al., 2014; Lankinen et al., 2022). Moreover, feedforward responses are not expected to be restricted exclusively to layer IV, as thalamocortical input rapidly propagates through canonical intracortical circuits to supragranular layers II/III (Callaway, 1998; Douglas & Martin, 1991). Consequently, a superficial-biased feedforward profile may better capture the laminar signature of feedforward-dominated processing measured at the spatiotemporal resolution of fMRI, which integrates both initial input and subsequent intracortical signal propagation. This provides a more physiologically plausible characterization of feedforward-related laminar profiles by accommodating both the known propagation of feedforward signals across cortical layers and residual measurement-related superficial biases.. In addition, an empirical feedforward template was derived from the mean normalized visual-motion laminar profile of sighted controls. This empirical template provides an independent validation of the expected feedforward laminar organization, as visual motion processing in hMT+/V5 of sighted individuals is established to rely on canonical ascending sensory pathways. All model profiles were normalized to the range [−1, 1] to enable correlation with observed participant layer profiles in permutation testing.

VASO-derived layer-wise estimates were extracted for each subject. Within each subject, laminar profiles were rescaled using a Min-Max normalization to the range [−1, 1] to remove inter-individual amplitude differences while preserving profile shape. To identify atypical laminar profiles, subjects showing consistent layer-wise deviations across multiple depths were flagged as potential outliers. Because inclusion of such profiles inflates group variance estimates, group-level standard deviations were computed after temporarily excluding flagged subjects, yielding a stable estimate of dispersion. Subjects whose layer-wise values exceeded ±3 standard deviations of this estimate were excluded from subsequent analyses (see Fig.S7). For each subject, Pearson correlations were computed between the observed laminar profile and each model template. Group-level observed correlations were obtained by averaging across subjects. Statistical significance was assessed using non-parametric permutation testing: for each subject, laminar labels were randomly shuffled 100,000 times, correlations recomputed, and null distributions generated. Empirical p-values were defined as the proportion of permutations yielding correlations greater than or equal to the observed group-level correlation.

#### Layer-wise activity statistical analyses against baseline

Statistical analyses were performed on layer-resolved fMRI signals extracted from anatomically and functionally defined regions of interest. Data consisted of subject-averaged parameter estimates derived from single-subject GLM contrasts and sampled across three equi-count cortical depth bins. All analyses were conducted in R (version 4.4.2; R Core Team, 2024) using custom scripts and standard statistical libraries. Analyses were performed separately for group, sensory modality, and region of interest. To assess whether activity within each cortical depth bin significantly deviated from baseline, we conducted layer-specific one-sample non-parametric tests using a one-sided Wilcoxon signed-rank test (activity > 0). P-values were computed separately for each cortical depth. To account for multiple comparisons across layers, we applied false discovery rate (FDR) correction using the Benjamini–Hochberg procedure.

## ACKNOWLEDGEMENTS

Computational resources have been provided by the supercomputing facilities of the Université Catholique de Louvain (CISM/UCL) and the Consortium des Équipements de Calcul Intensif en Fédération Wallonie Bruxelles (CÉCI) funded by the Fond de la Recherche Scientifique de Belgique (F.R.S.-FNRS) under convention 2.5020.11 and by the Walloon Region. We thank Dr. Ceren Battal and Dr. Mohamed Rezk for insightful discussions which provided important conceptual and methodological foundations for the present study. f/MRI data were acquired at the GIGA-Cyclotron-In Vivo Imaging Research Centre at the University of Liège (Belgium). We thank Dr Laurent Lamalle and Dr. Siya Sherif for the assistance in the early phases of data acquisition. We thank Rüdiger Stirnberg (DZNE, Bonn) for providing the sequence code used for acquiring VASO data. We also thank Dr. Peter Bandettini (FMRIF Chief at NIMH) for providing the support and resources that allowed us to start the preprocessing of the sub-millimetre fMRI data and Dr. Paul A. Taylor (SSCC Director at NIMH) for the help provided for the implementation of AFNI software in this study. OC is a senior research associate at the Fond National de la Recherche Scientifique de Belgique (FRS-FNRS).

## FUNDING

The project was funded in parts by the Belgian Excellence of Science (EOS) program (Project No. 30991544) awarded to OC; a Flagship ERA-NET grant SoundSight (FRS-FNRS PINT-MULTI R.8008.19) awarded to OC, and a PDR F.R.S-FNRS grant (BlindLayer) attributed to MB, JM and OC. RH is currently supported by the grant 5P41EB030006-05 (PI: Rosen).

## AUTHOR CONTRIBUTION

Conception and design: MB, RH, JM, OC. Administrative support: MB, RG, OC; Provision of study materials or patients: MB, EG, RH, RG, MVB, OC; Collected data: MB, EG, MVB. Analysed data: MB, with supervision support from RH and OC. Wrote the original draft: MB, OC. Writing - Review & Editing: All authors. Main funding acquisition: OC.

## COMPETING INTERESTS

The authors declare no competing interests.

## DATA AVAILABILITY STATEMENT

Raw f/MRI data are not publicly available as full anonymity of the participants cannot be guaranteed, even after defacing the MRI images and due to the lack of explicit consent from our participants. Code used to process the data and processed data to replicate statistical results and figures can be found on GitHub: https://github.com/marcobarilari/analysis_high-res_2023_Vaso_Blind_Layer_Motion

## SUPPLEMENTARY MATERIAL

**Method S1. Behavioral performance during the motion localizers.**

To ensure participants’ attention during the visual and auditory motion localizers, sighted controls (SC) were instructed to detect a fixation cross color change (from red to green) in both localizers. During the auditory localizer, both SC and early blind participants (EB) were additionally asked to detect a target sound that either moved faster during moving blocks or was shorter during static blocks. Behavioral data from two early blind participants were not recorded due to button-box malfunction, and on SC participants did not perform the fixation task during the auditory localizer; these participants were excluded from the behavioral analyses. Overall, participants were highly accurate in both tasks. Mean accuracy for the sound target was high in both groups, with EB participants achieving 98% (SD = 2.8%) and SC participants 93.4% (SD = 8.8%), see Fig.S2a. A Wilcoxon rank-sum test indicated no significant difference between groups, *W* = 43, *p* = 0.23, confirming comparable performance. SC participants performed near ceiling on the fixation task during both localizers (auditory: mean = 98.6%, SD = 3.2%; visual: mean = 99.7%, SD = 1.1%), see Fig.S2b. A paired Wilcoxon signed-rank test showed no significant difference in fixation performance between the visual and auditory localizers, *V* = 6, *p* = 0.181, indicating consistent attention across modalities.

**Method S2. Behavioral performance during the VASO layer-profile experiment,**

During the main VASO experiment, participants performed a one-back task adapted to stimulus modality (visual or auditory) and stimulation type (static: dots changing color and sound changing noise or moving stimuli: same direction repeated twice in the horizontal or vertical axes). Sighted controls (SC) also performed a fixation task during the visual version of the experiment (see Methods). In the auditory experiment, both early blind (EB) and sighted control (SC) participants performed the one-back task. In the visual experiment, only SC participants were tested. Behavioral accuracy was computed as the proportion of correct responses per subject and condition. For the auditory task, a mixed-design ANOVA with group (EB, SC) as a between-subject factor and measure (static vs motion) as a within-subject factor was performed. For the visual task (SC only), paired Wilcoxon signed-rank tests were used to compare static vs motion and horizontal vs vertical conditions due to ceiling performance and non-normal distributions. In the auditory one-back task, accuracy was high during static blocks in both groups (EB: mean = 98.4%, SD = 3.3%; SC: mean = 98.6%, SD = 2.3%) but decreased during motion blocks (EB: mean = 86.2%, SD = 15.3%; SC: mean = 71.0%, SD = 18.7%), see Fig.S3a. A mixed-design ANOVA with group (EB, SC) as a between-subject factor and condition (static, motion) as a within-subject factor revealed no significant main effect of group, *F*(1,16) = 3.12, *p* = 0.1, but a significant main effect of condition, *F*(1,16) = 20.01, *p* < 0.001, reflecting lower accuracy during motion compared to static stimulation The group × condition interaction showed no difference between the groups, *F*(1,16) = 2.8, *p* = 0.11 (Fig.S3a).

When separating motion direction (Fig.S3d), performance was lower for vertical compared to horizontal motion in both groups (EB: horizontal mean = 98.3%, SD = 2.6%; vertical mean = 74.07%, SD = 31.4%; SC: horizontal mean = 83.1%, SD = 15.9%; vertical mean = 59.0%, SD = 30.3%), consistent with the overall motion-related accuracy decrease. A mixed-design ANOVA with group (EB, SC) as a between-subject factor and condition (horizontal, vertical) as a within-subject factor revealed no significant main effect of group, F(1,16) = 3.53, p = 0.08, but a significant main effect of condition, F(1,16) = 10.48, p < 0.005, reflecting lower accuracy during vertical compared to horizontal stimulation. The group × condition interaction showed no difference between the groups, F(1,16) = 0, p = 0.99 (Fig.S3d). In the visual one-back task (SC only; Fig.S3b), performance was near ceiling in both static (mean = 98.8%, SD = 3.0%) and motion conditions (mean = 97.2%, SD = 4.0%). A paired Wilcoxon signed-rank test revealed no difference between conditions, *V* = 10, *p* = 0.09. No significant difference was observed between horizontal (M = 96.3%, SD = 6.0%) and vertical motion (M = 98.1%, SD = 3.0%), *V* = 2, *p* = 0.345 (Fig.S3e). Fixation task performance during the visual experiment remained stable across conditions (static: M = 94.1%, SD = 10.8%; motion: M = 94.1%, SD = 17.7%), with no significant difference, *V* = 8, *p* = 0.672 (Fig.S3c). Similarly, fixation accuracy did not differ between horizontal and vertical blocks (horizontal: 93.8%, SD = 17.7%; vertical: 94.4%, SD = 17.8%), *V* = 0, *p* = 1 (Fig. S3f). Overall, these results indicate high task compliance across experiments. In the auditory task, motion stimulation induced a measurable decrease in accuracy, whereas performance in the visual task remained near ceiling, suggesting that differences in laminar responses are unlikely to be driven by gross performance disparities.

**Method S3. Regions of interest cortical thickness control analyses.**

Cortical thickness values were extracted as a by-product of the laminar segmentation procedure implemented in LayNii (LN2_LAYERS, v2.6.0; L. Huber et al., 2024; L. (Renzo) Huber et al., 2021) via the layer-MAP (v0.2.0; https://github.com/marcobarilari/layer-MAP; Barilari et al., 2026). Thickness was computed within each individually defined fROI for each participant and computed from the equi-count laminar segmentation output within the fROI. To ensure adequate laminar sampling given the functional resolution (0.75 mm isotropic), we quantified mean cortical thickness in all fROIs. Across groups and regions, mean thickness was approximately 2.8 mm (Fig.S12), allowing at least one voxel to sample each of the deep, middle, and superficial compartments. To rule out potential structural confounds that could bias depth-dependent functional analyses, we statistically compared cortical thickness across regions and groups. Within sighted controls (SC), cortical thickness in right hMTa and right hMT+/V5 was compared using paired two-tailed t-tests. Between-group comparisons (SC right hMTa vs EB right hMT+/V5; SC right hMT+/V5 vs EB right hMT+/V5) were performed using independent two-sample two-tailed t-tests. No significant differences were observed across any comparison (all p > 0.11, p-adjusted for multiple comparisons using the Benjamini-Hochberg method: all p > 0.33).

**Figure S1.**
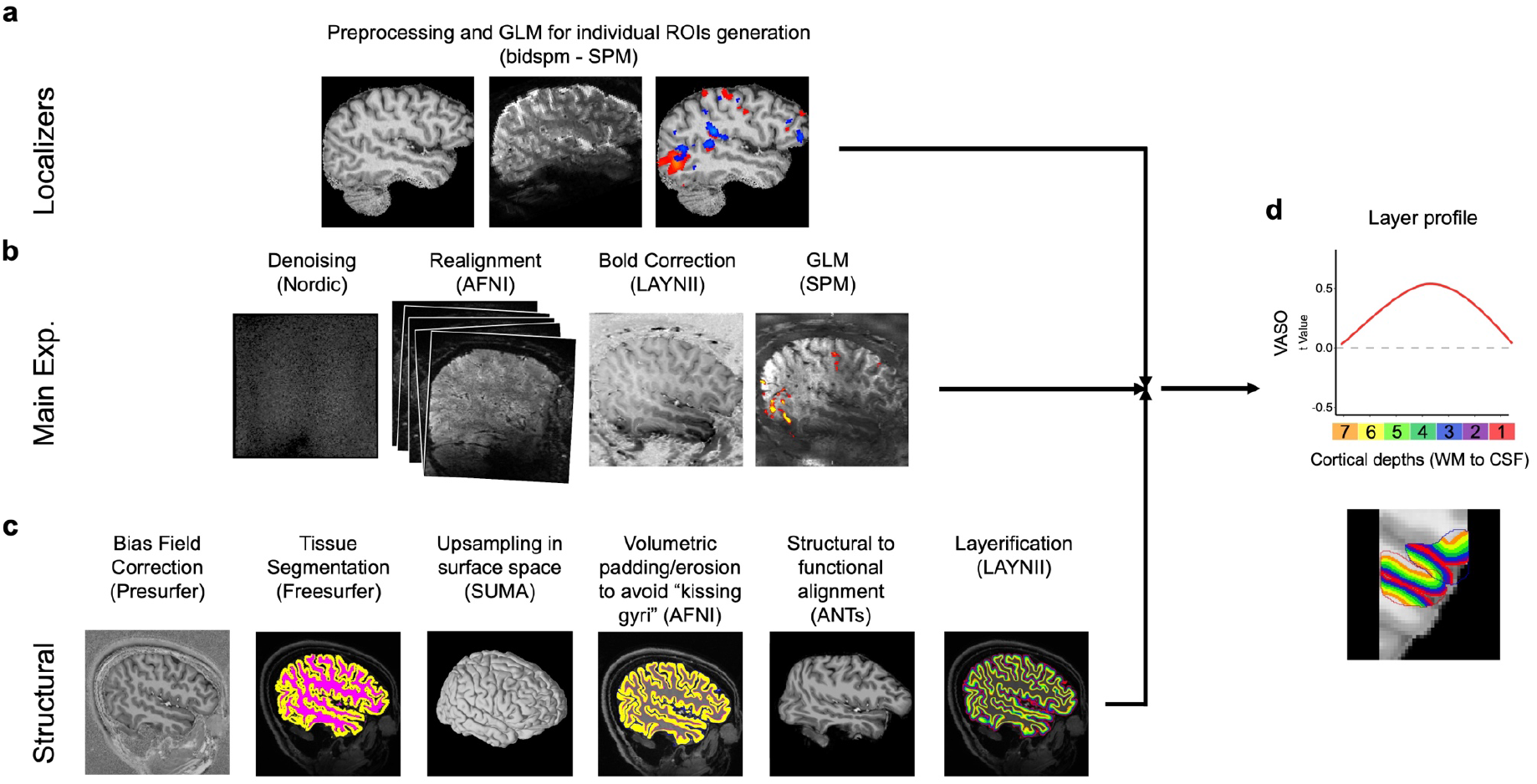
Functional and structural schematic of the analyses pipeline. In order to obtain a functional layer profile from each subject, **a,** whole brain functional data (1.6 mm isotropic) where pre-processed and statistically analysed using SPM via bidspm. **b,** Sub-millimetre SS-SI VASO functional data (0.75 mm isotropic) where analysed using a custom pipeline implementing different software depending on the analyses step: NORDIC for data denoising, AFNI for spatial realignment, BOLD correction of the VASO data using LayNii, SPM for statistical analyses. **c,** Anatomical MP2RAGE images (0.75 mm isotropic) where processed using the layerfMRI toolbox implementing different software depending on the analyses step: Presurfer with SPM for bias field correction, Tissue segmentation using Freesurfer, up-sampling in surface space using SUMA, Volumetric padding/erosion using AFNI, structural alignment distortion to match the functional space using ANTs, and cortical depth bins “layerification” using LayNii. **d,** the output of these three pipelines is pulled together to obtain the layer activity profile from within a specific region of interest (fROI).

**Figure S2.**
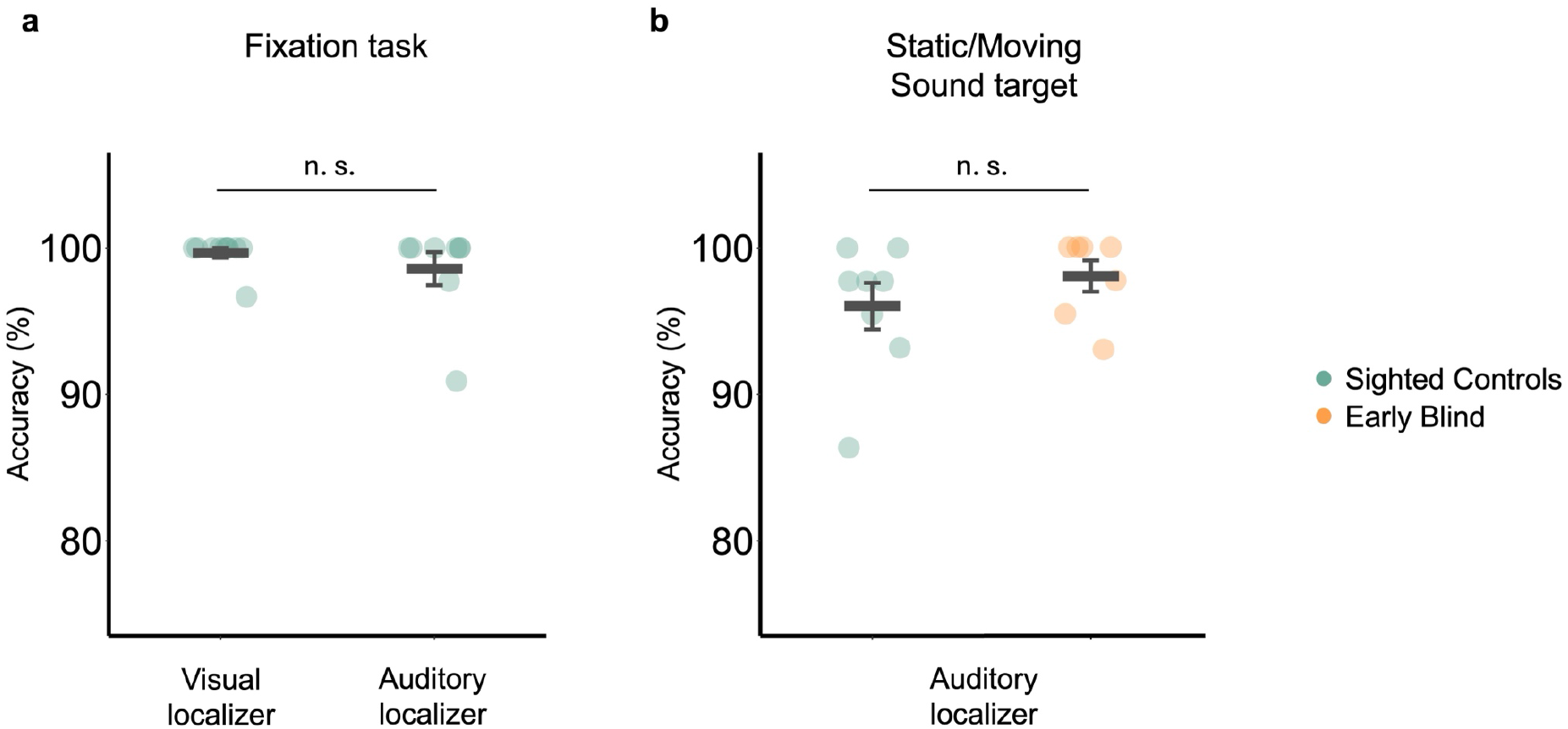
Behavioural performance during the motion localizers. a, Fixation detection accuracy in sighted control (SC) participants during the visual and auditory motion localizers. Points represent individual subjects divided by localizer type, bars indicate group means, and error bars represent the standard error. Performance was near ceiling in both localizers, with no significant difference between tasks. **b,** Sound target detection accuracy during the auditory motion localizer for early blind (EB) and sighted control (SC) participants. Points represent individual subjects, bars indicate group means, and error bars represent the standard error. No significant group differences were observed.

**Figure S3.**
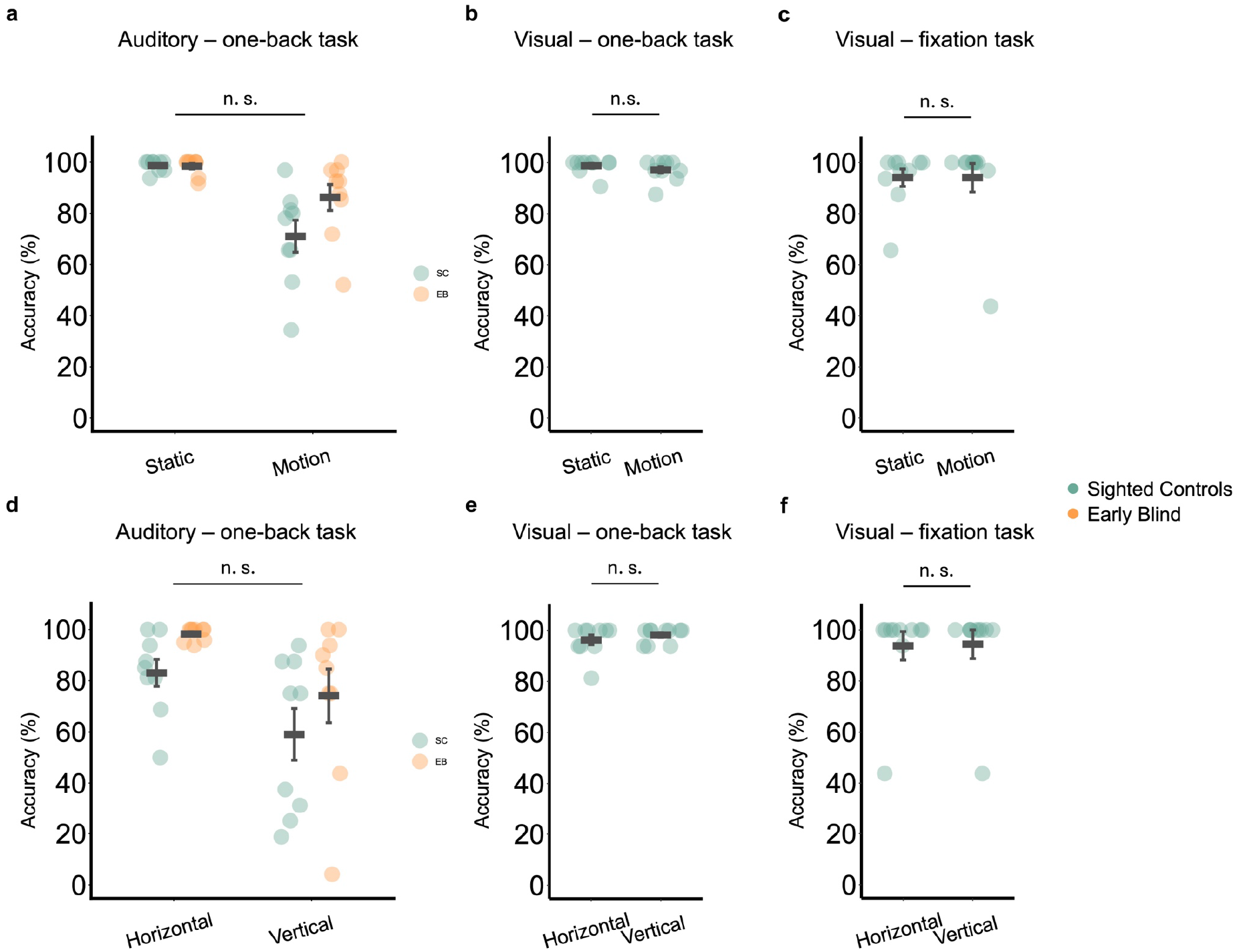
Behavioral performance during the VASO experiment. a,b,c, Accuracy for static and motion conditions. **a,** Auditory one-back task (EB, early blind; SC, sighted controls) for the static and motion conditions. A mixed-design ANOVA revealed a main effect of condition (*F*(1,16) = 20.01, *P* < 0.001), no main effect of group (*F*(1,16) = 3.12, *P* = 0.1), and no significant group × condition interaction (*F*(1,16) = 2.8, *P* = 0.11). **b,** Visual one-back task (SC only) showed a small but significant difference between static and motion conditions (Wilcoxon signed-rank test, *V* = 167, *P* < 0.001). c, Visual fixation task (SC only) showed no difference between conditions (*V* = 8, *P* = 0.67). **d,e,f,** Accuracy for horizontal and vertical conditions for the same tasks as in **a,b,c. d,** Auditory one-back task (EB, early blind; SC, sighted controls) for the horizontal and vertical conditions. A mixed-design ANOVA revealed no significant main effect of group (F(1,16) = 3.53, p = 0.08), a significant main effect of condition (F(1,16) = 10.48, p < 0.005), reflecting lower accuracy during vertical compared to horizontal stimulation, and no significant group × condition interaction (F(1,16) = 0, p = 0.99). **e,** Visual one-back task showed no significant directional difference (*V* = 2, *P* = 0.35). **f,** Visual fixation task showed no difference (*V* = 0, *P* = 1.00). Points indicate individual participants; bars show mean ± standard error.

**Figure S4.**
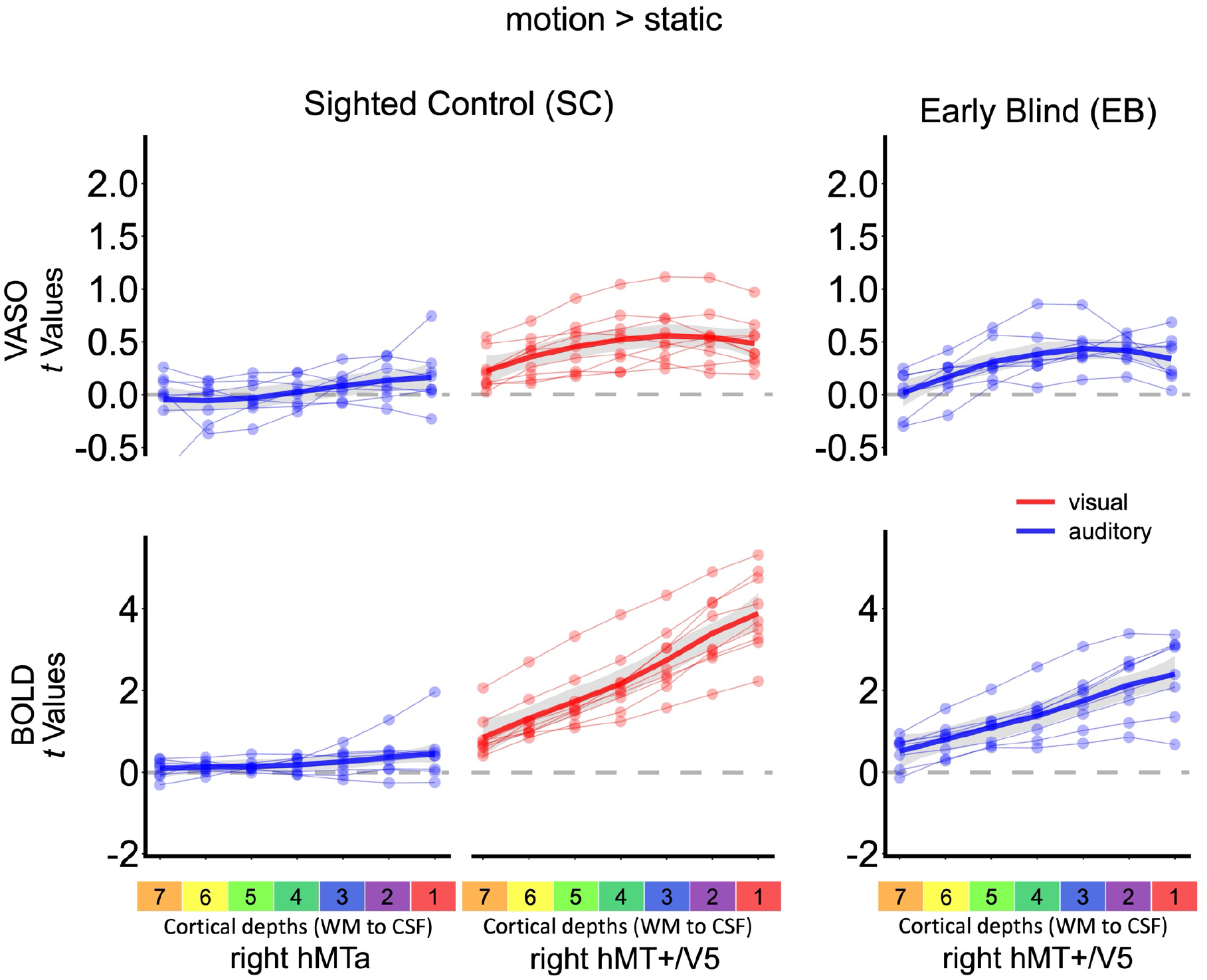
Motion > Static laminar activity profiles for VASO and BOLD. Related to. Fig. 3c. Activation layer profiles for the [motion > static] contrast are shown for each region of interest (fROI), group and modality. Results are presented as *t*-values from the GLM for VASO (top rows) and BOLD (bottom rows). Thin lines represent individual participants, with data points connected across cortical depth to illustrate single-subject laminar profiles; thick lines indicate the group mean.

**Figure S5.**
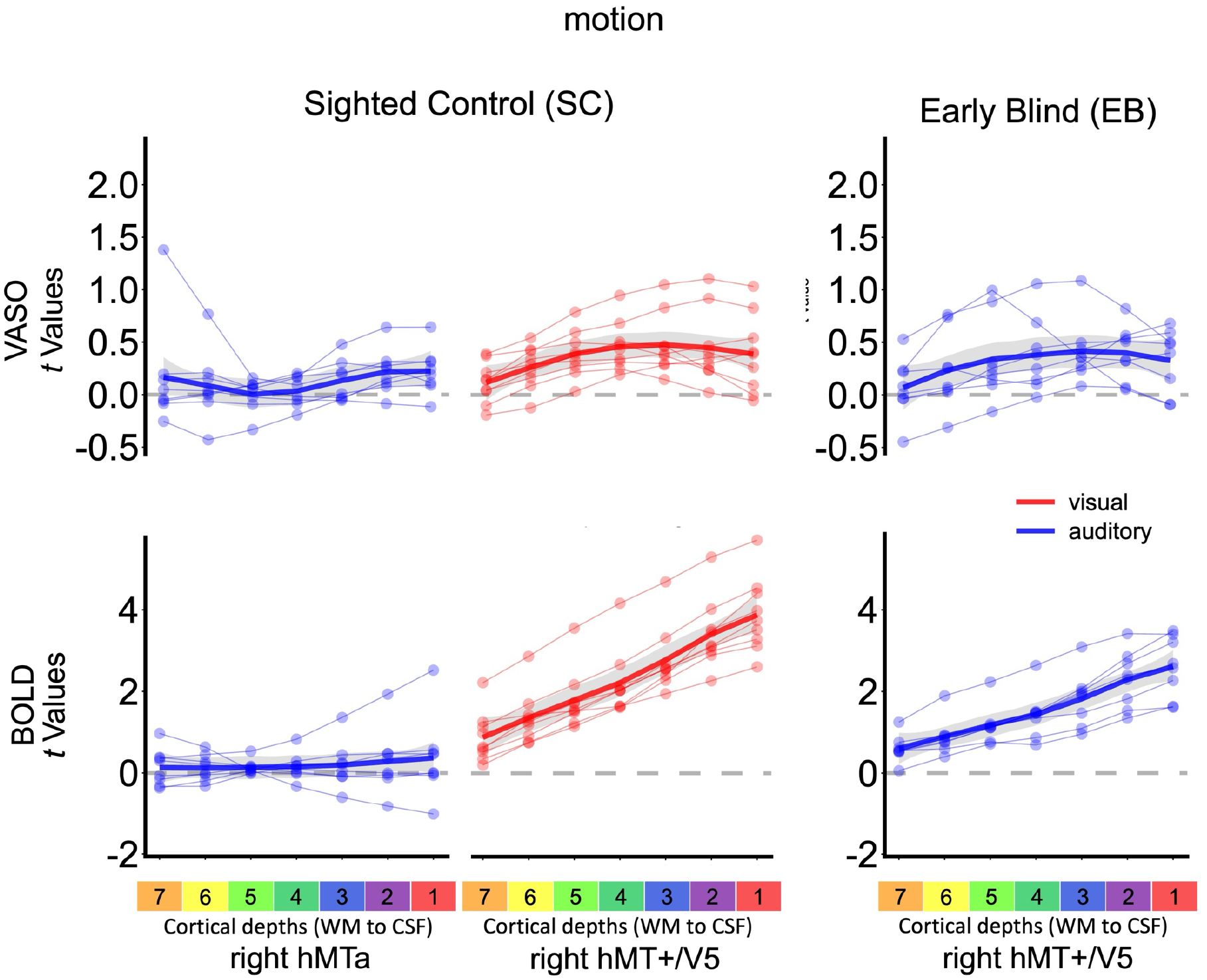
Motion laminar activity profiles for VASO and BOLD. Related to. Fig. 3c. Activation layer profiles for the [motion] condition are shown for each region of interest (fROI), group and modality. Results are presented as *t*-values from the GLM for VASO (top rows) and BOLD (bottom rows). Thin lines represent individual participants, with data points connected across cortical depth; thick lines indicate the group mean.

**Figure S6.**
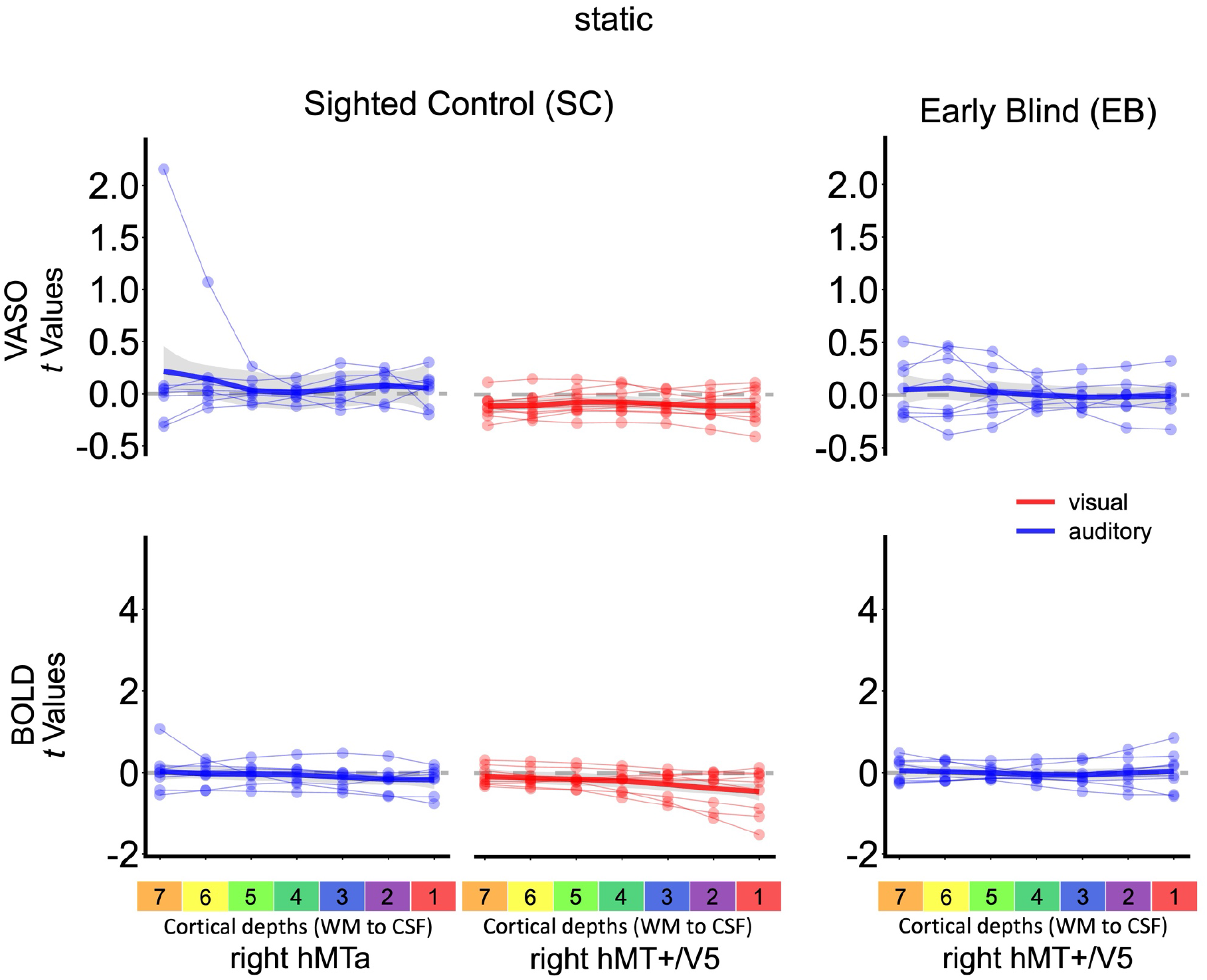
Static laminar activity profiles for VASO and BOLD. Related to. Fig. 3c. Activation layer profiles for the [static] condition are shown for each region of interest (fROI), group and modality. Results are presented as *t*-values from the GLM for VASO (top rows) and BOLD (bottom rows). Thin lines represent individual participants, with data points connected across cortical depth; thick lines indicate the group mean.

**Figure S7.**
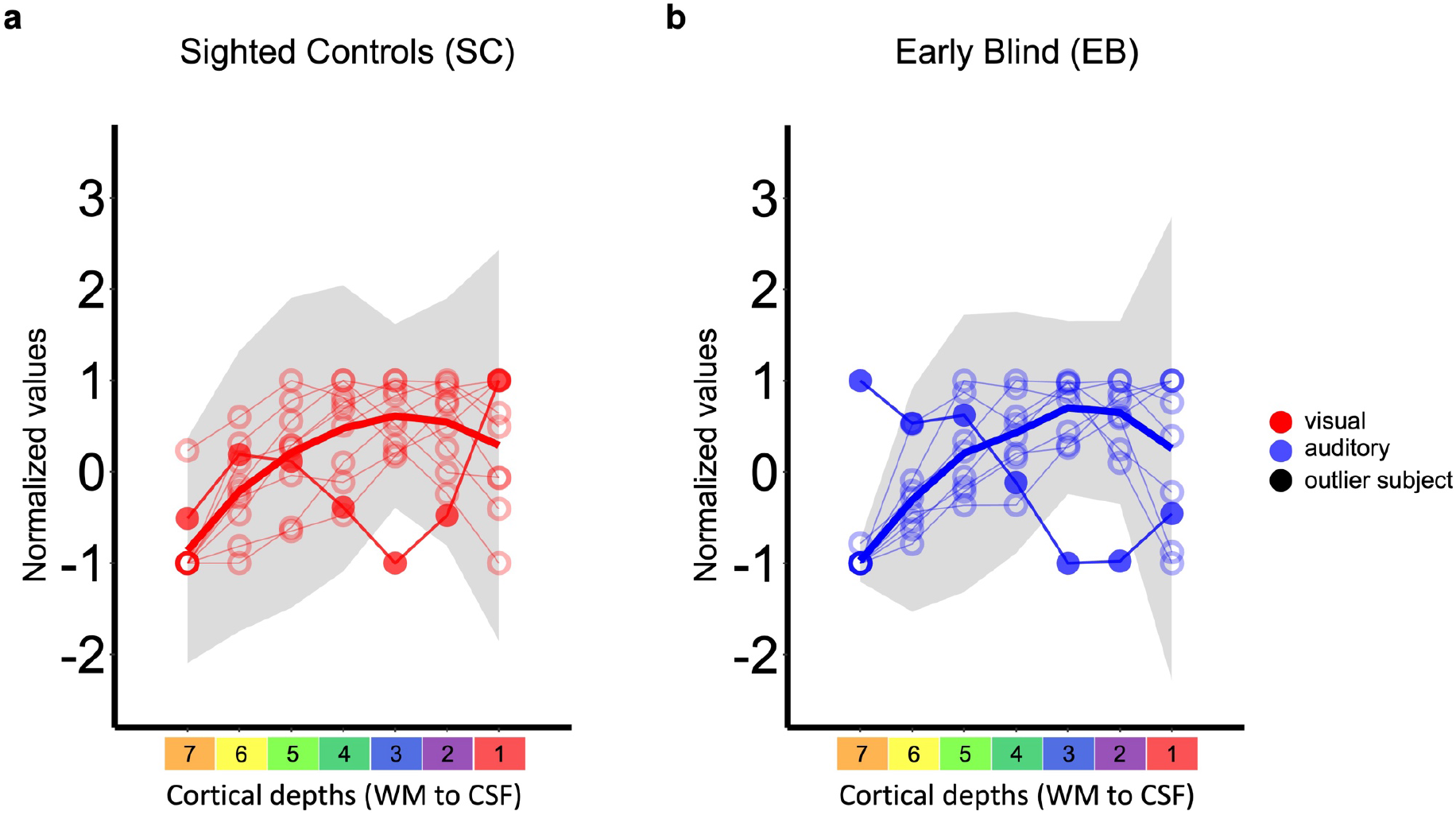
Normalized cortical motion selectivity profile and criteria for exclusion for the model comparison analyses. Related to Fig.3d. Cortical selectivity is normalized using a Min-Max approach within each subject result for the right hMT+/V5 in **a,** sighted for the visual and in **b,** early blind for the auditory modality (right). Light grey shade depicts the +/- 3 standard deviation (SD) for each cortical depth computed excluding a priori the deviant subjects (one sighted and one early blind). Filled datapoints represent the outliers showing that for some layers it lais outside the 3 SD, not-filled dots represent the subjects’ datapoint within the 3 SD.

**Figure S8.**
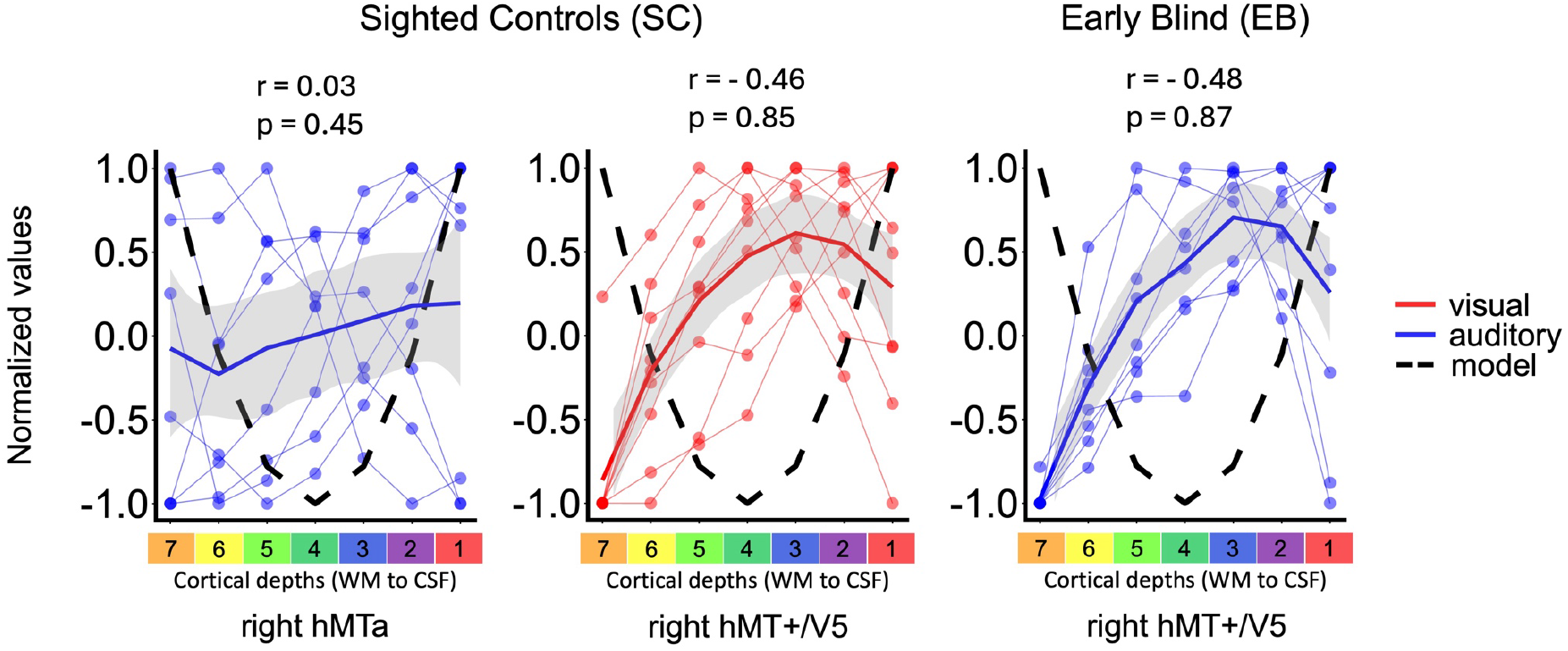
Permutation-based inference for the feedback laminar profile model (“U-shaped”). Related to Fig.3d. Permutation test assessing the correspondence between subject-specific VASO-derived laminar response profiles (light-colored lines and dots) and the feedback (“U-shaped”, dashed black line) model across seven equi-count cortical depths across group, region of interest (fROI), and sensory modality. For each subject, depth-wise responses were normalized to the range [−1, 1] and correlated with the predefined feedback model template (dashed line). Null distributions were generated by randomly shuffling laminar depth labels within subjects (100,000 permutations) and recomputing correlations. Empirical p-values correspond to the proportion of permuted correlations equal to or greater than the observed group-level correlation. Solid-colored lines represent the group mean layer profile.

**Figure S9.**
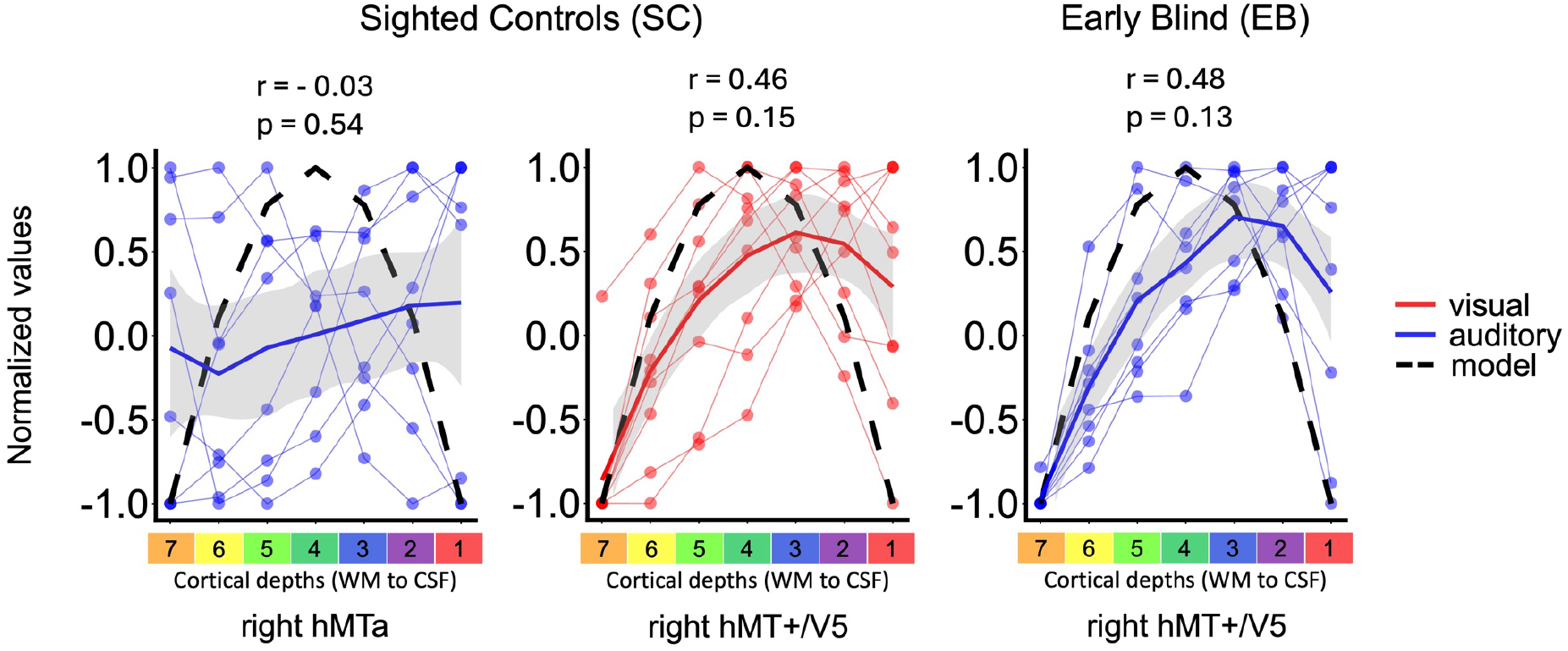
Permutation-based inference for the symmetrical feedforward laminar profile model (“inverted-U-shaped). Related to Fig.3d. Permutation test assessing the correspondence between subject-specific VASO-derived laminar response profiles (light-colored lines and dots) and the canonical feedforward (“inverted-U-shaped”, dashed black line) model across seven equi-count cortical depths across group, region of interest (fROI), and sensory modality. For each subject, depth-wise responses were normalized to the range [−1, 1] and correlated with the predefined feedback model template. Null distributions were generated by randomly shuffling laminar depth labels within subjects (100,000 permutations) and recomputing correlations. Empirical p-values correspond to the proportion of permuted correlations equal to or greater than the observed group-level correlation. Solid-colored lines represent the group mean layer profile.

**Figure S10.**
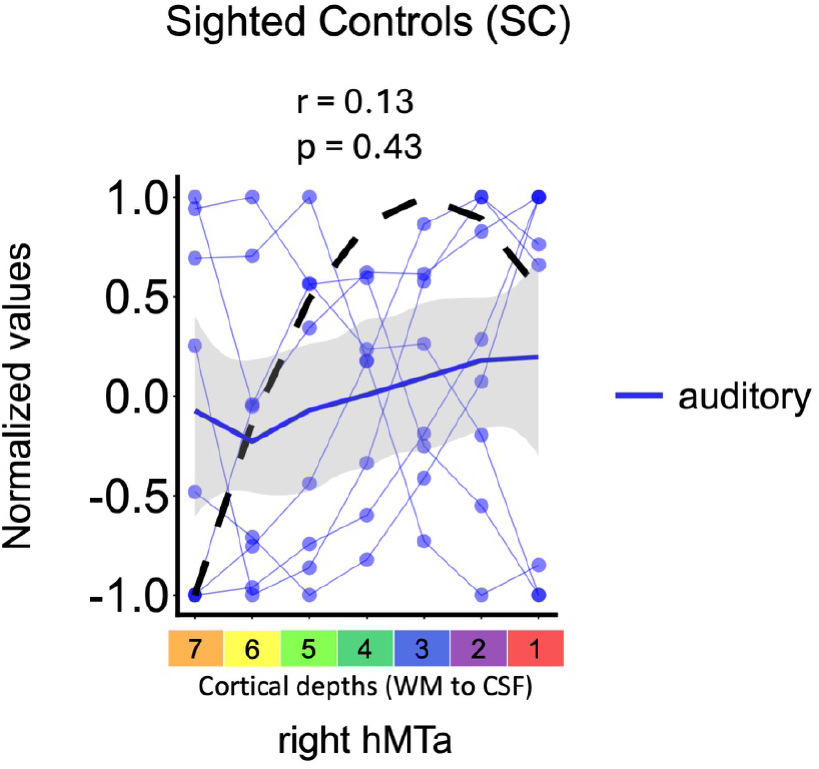
Permutation-based inference for the skewed feedforward laminar profile model (“skewed inverted-U-shaped) for the auditory modality in the right hMTa. Related to. Fig.3 Permutation test assessing the correspondence between subject-specific VASO-derived laminar response profiles (light-blue lines and dots) and the skewed feedforward (“inverted-U-shaped”, dashed black line) model across seven equi-count cortical depths across group, region of interest (fROI). For each subject, depth-wise responses were normalized to the range [−1, 1] and correlated with the predefined feedback model template. Null distributions were generated by randomly shuffling laminar depth labels within subjects (100,000 permutations) and recomputing correlations. Empirical p-values correspond to the proportion of permuted correlations equal to or greater than the observed group-level correlation. Solid-colored line represents the group mean layer profile.

**Figure S11.**
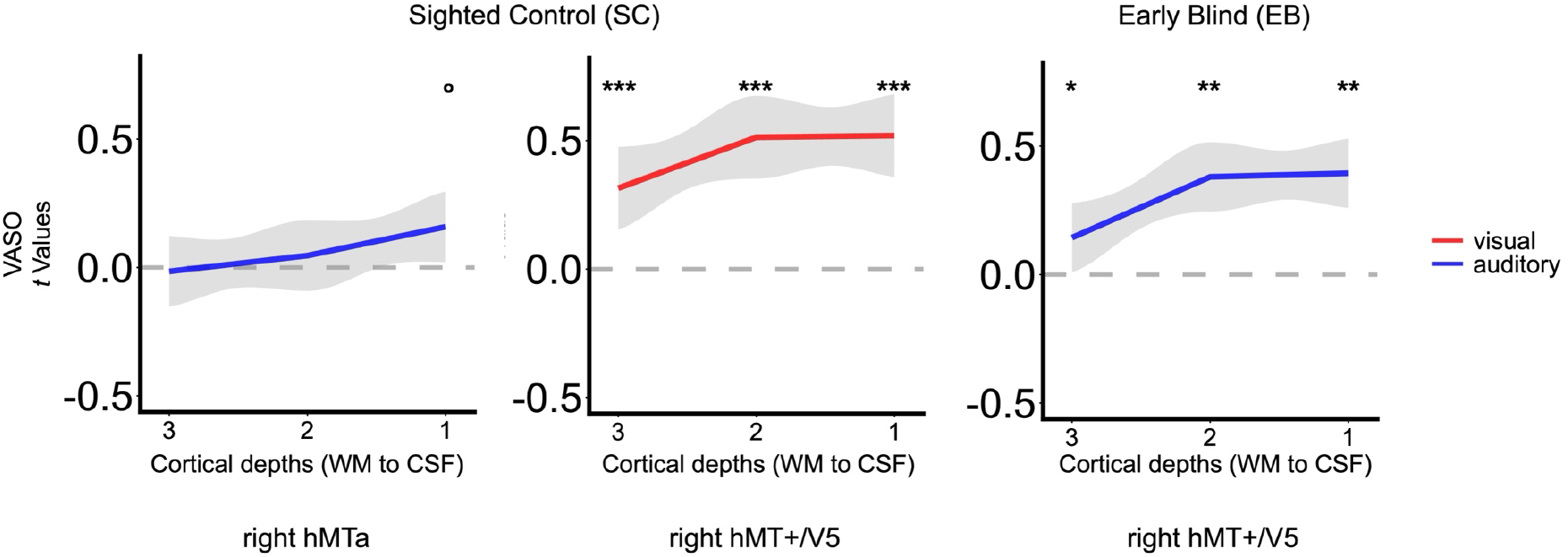
Layer-wise activity relative to baseline. Layer-resolved fMRI responses are shown for each cortical depth bin (three equi-count layers) within anatomically and functionally defined regions of interest, separately for group, region of interest (fROI), and sensory modality. Values represent subject-averaged parameter estimates extracted from single-subject GLM contrasts. Statistical significance against baseline was assessed independently at each depth using one-sided Wilcoxon signed-rank tests (activity > 0), with false discovery rate (FDR) correction applied across layers. * p < 0.5; ** p < 0.1; *** p < 0.001; ° p-value significant not corrected for multiple comparison.

**Figure S12.**
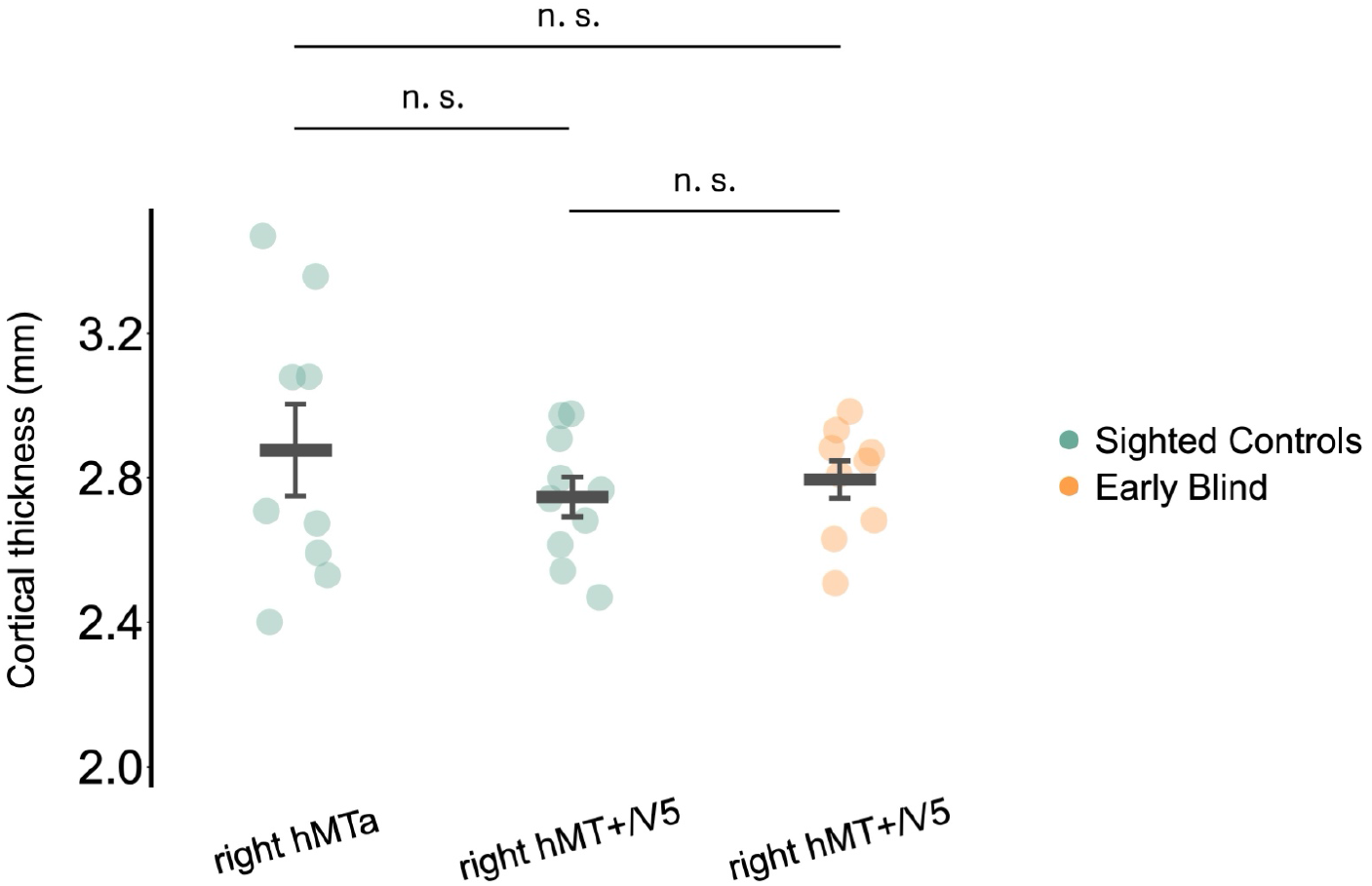
Cortical thickness across the regions of interest and groups. Mean cortical thickness (mm) measured within each individually defined fROI (hMTa in sighted controls; hMT+/V5 in sighted controls and early blind participants). Thickness values were obtained as a by-product of the laminar segmentation procedure (LN2_LAYERS, LayNii). Horizontal bar represents group mean and error bars indicate ± standard error. Across fROIs, mean cortical thickness was approximately 2.8 mm, allowing at least one 0.75 mm isotropic voxel to sample each of the deep, middle, and superficial depth compartments. No significant differences in cortical thickness were observed across fROI or group comparisons (all p > 0.11, p-adjusted for multiple comparison using Benjamini-Hochberg method: all p > 0.33; see Method S3).

**Table S1.** Demographic information of the Sighted Control (SC) and Early Blind (EB) participants. Group: EB = Early Blind, SC = Sighed Controls; age and education are expressed in years; sex: F = female, M = male. Blindness cause: RFO = retrolental fibroplasia from neonatal over-oxygenation; LCA = Leber’s congenital amaurosis.

| Subject | Group | Age | Sex | Education | Blindness onset | Blindness Cause | Light Perception |
| --- | --- | --- | --- | --- | --- | --- | --- |
| EB01 | EB | 52 | M | 15 | 0 | RFO | Yes, left eye |
| EB02 | EB | 36 | M | 15 | 0 | LCA | Yes |
| EB03 | EB | 45 | F | 15 | 0 | LCA | No |
| EB05 | EB | 25 | F | 12 | 2 | Retinoblastoma | Yes |
| EB06 | EB | 27 | M | 15 | 0 | LCA | No |
| EB07 | EB | 33 | M | 12 | 0 | LCA | No |
| EB09 | EB | 48 | M | 12 | 0 | LCA | Yes |
| EB10 | EB | 44 | M | 17 | 0 | LCA | No |
| EB11 | EB | 37 | M | 12 | 0 | LCA | No |
| SC03 | SC | 54 | M | 15 | - | - | - |
| SC04 | SC | 29 | M | 15 | - | - | - |
| SC07 | SC | 50 | M | 17 | - | - | - |
| SC08 | SC | 24 | F | 15 | - | - | - |
| SC09 | SC | 57 | M | 15 | - | - | - |
| SC010 | SC | 26 | F | 17 | - | - | - |
| SC011 | SC | 30 | F | 17 | - | - | - |
| SC012 | SC | 24 | M | 15 | - | - | - |
| SC013 | SC | 38 | F | 21 | - | - | - |
| SC014 | SC | 31 | M | 21 | - | - | - |

**Table S2.** Group results in MNI space of the univariate analyses for the visual and auditory motion localizer in sighted controls and early blind. Related to Fig.2a. Results for the contrast [motion>static] are corrected for multiple comparisons using family-wise error correction (FWE) across the whole brain at a significance level of p<0.05 or over small volume correction (sphere of 10 mm). Coordinates used for small volume correction were extracted by looking at the map at p = 0.001 uncorrected and extracting the sphere center from the local global maxima of a cluster of activation. Coordinates are as follow: *: left Middle Temporal Gyrus (L hMT+/V5) [-40 −7-12] and right Middle Temporal Gyrus (R hMT+/V5)[46 −64 10] from the SC visual [motion > static] results; ** Left Superior Temporal Gyrus (L Planum Temporale) [-53 −28 8] from the SC auditory [motion>static] results; *** Left Cuneus [-10 −84 36] and right Cuneus [13 −78 39] from the EB auditory [motion>static] results. Coordinates are reported in MNI space. L: Left; R: Right; G: Gyrus.

| Area | Cluster size (k) | x (mm) | y (mm) | z (mm) | z | t | p(FEW) |
| --- | --- | --- | --- | --- | --- | --- | --- |
| SC visual [motion > static] |  |  |  |  |  |  |  |
| L Superior Occipital | 527 | -17 | -85 | 13 | 6.57 | 24.97 | < 0.001 |
| L Lingual G | 1017 | -6 | -80 | 0 | 5.74 | 15.28 | 0.002 |
| R Middle Occipital G | 32 | 28 | -88 | 16 | 5.21 | 11.47 | 0.021 |
| L Lingual G | 6 | -24 | -67 | -6 | 5.14 | 11 | 0.028 |
| L Lingual G | 13 | -17 | -86 | -9 | 5.13 | 10.99 | 0.028 |
| L Lingual G | 20 | -11 | -81 | -11 | 5.09 | 10.77 | 0.033 |
| L Middle Temporal G<br>(L hMT+/V5) | 900 | -38 | -62 | 10 | 4.54 | 8.11 | 0.001* |
| R Middle Temporal G<br>(R hMT+/V5) | 997 | 39 | -59 | 5 | 4.38 | 7.53 | 0.001* |
| SC auditory [motion > static] |  |  |  |  |  |  |  |
| R Superior Temporal G<br>(R Planum Temporale) | 19 | 65 | -22 | 2 | 5.29 | 11.92 | 0.015 |
| L Superior Temporal G<br>(L Planum Temporale) | 715 | -53 | -28 | 8 | 4.82 | 9.38 | < 0.001** |
| R Middle Temporal G<br>(R hMT+/V5) | 88 | 49 | -56 | 8 | 3.01 | 3.86 | 0.05* |
| EB auditory motion > static] |  |  |  |  |  |  |  |
| R Superior Temporal G<br>(R Planum Temporale) | 905 | 44 | -28 | 12 | 4.47 | 9.11 | 0.001*** |
| L Superior Temporal G<br>(L Planum Temporale) | 681 | -38 | -30 | 7 | 4.36 | 8.55 | 0.002*** |
| L Cuneus | 534 | -9 | -81 | 36 | 4 | 7.01 | 0.005*** |
| R Middle Temporal G<br>(R hMT+/V5) | 518 | 46 | -65 | 8 | 3.99 | 6.97 | 0.005* |
| L Middle Temporal G<br>(L hMT+/V5) | 282 | -43 | -65 | 8 | 3.69 | 5.92 | 0.012* |
| R Cuneus | 66 | 9 | -85 | 44 | 3.23 | 4.64 | 0.037*** |
| EB > SC [motion > static] |  |  |  |  |  |  |  |
| L Cuneus | 261 | -11 | -81 | 36 | 3.99 | 5.02 | 0.003*** |
| R Cuneus | 518 | 10 | -83 | 44 | 3.91 | 4.88 | 0.004*** |
| R Middle Temporal G<br>(R hMT+/V5) | 126 | 46 | -57 | 7 | 3.75 | 4.59 | 0.006* |
| L Middle Temporal G<br>(L hMT+/V5) | 25 | -40 | -64 | 5 | 3.49 | 4.17 | 0.013* |

**Table S3.** Peak coordinates of individually defined motion-selective regions of interest in MNI space. Related to Fig.2b. Peak coordinates for hMT+/V5 and hMTa were identified at the single-subject level from the contrast *[motion > static]* using the appropriate modality-specific motion localizer (visual motion localizer: hMT+/V5 in sighted controls; auditory motion localizer: hMT+/V5 in early blind participants and hMTa in sighted controls). EB, early blind; SC, sighted controls; L, left hemisphere; R, right hemisphere.

| Subject | Group | R hMTa |  |  | R hMT+/V5 |  |  |
| --- | --- | --- | --- | --- | --- | --- | --- |
|  |  | x (mm) | y (mm) | z (mm) | x (mm) | y (mm) | z (mm) |
| EB01 | EB | -- | -- | -- | 46 | -64 | 5 |
| EB02 | EB | -- | -- | -- | 44 | -70 | 8 |
| EB03 | EB | -- | -- | -- | 59 | -64 | 13 |
| EB05 | EB | -- | -- | -- | 44 | -77 | 0 |
| EB06 | EB | -- | -- | -- | 47 | -75 | -3 |
| EB07 | EB | -- | -- | -- | 44 | -59 | 2 |
| EB09 | EB | -- | -- | -- | 46 | -65 | 4 |
| EB10 | EB | -- | -- | -- | 39 | -57 | 12 |
| EB11 | EB | -- | -- | -- | 49 | -65 | -15 |
| SC03 | SC | -- | -- | -- | 39 | -65 | 10 |
| SC04 | SC | 46 | -57 | 10 | 46 | -67 | 5 |
| SC07 | SC | 41 | -60 | 4 | 51 | -68 | 7 |
| SC08 | SC | 60 | -56 | 15 | 46 | -64 | 8 |
| SC09 | SC | 51 | -57 | 7 | 44 | -65 | 7 |
| SC10 | SC | 51 | -65 | 12 | 47 | -62 | 5 |
| SC11 | SC | 52 | -72 | 13 | 49 | -67 | 7 |
| SC12 | SC | 49 | -60 | 10 | 43 | -64 | 0 |
| SC13 | SC | 44 | -70 | 20 | 41 | -73 | 8 |
| SC14 | SC | 46 | -56 | 5 | 43 | -68 | 7 |

**Table S4.** Statistical summary of layer-wise activity relative to baseline. Related to Fig.S11. P-values from layer-specific one-sample tests (activity > 0) corresponding to the results shown in Fig. S5. Tests were performed independently for each cortical depth bin, group, modality, and region of interest. Both uncorrected and corrected p-values are reported. Significance thresholds are indicated as * p < 0.05; ** p < 0.01; *** p < 0.001; ° p-value significant not corrected for multiple comparison.

| Group | fROI | Task | Cortical Depth | V | p-value (FDR) |
| --- | --- | --- | --- | --- | --- |
| SC | right hMTa | auditory<br>[motion > static] | 3 | 20 | 0.42 |
|  |  |  | 2 | 26 | 0.23 |
|  |  |  | 1 | 31 | 0.039 ° |
| SC | right hMT+/V5 | visual<br>[motion > static] | 3 | 55 | < 0.001 *** |
|  |  |  | 2 | 55 | < 0.001 *** |
|  |  |  | 1 | 55 | < 0.001 *** |
| EB | right hMT+/V5 | auditory<br>[motion > static] | 3 | 40 | 0.019 * |
|  |  |  | 2 | 45 | 0.003 ** |
|  |  |  | 1 | 45 | 0.003 ** |

**Table S5.** Psychophysiological interactions (PPI) from the visual and the auditory motion localizers in sighted controls and early blind individuals. *Related to Fig.2*. Results for the visual motion localizer contrast (visual [motion>static]) and the auditory motion localizer contrast (auditory [motion>static]). Seed regions comprised individually defined right visual-defined hMT+/V5 for sighted controls in the visual localizer PPI and right auditory-defined hMT+/V5 for both groups in the auditory localizers group comparison. * The coordinate for small-volume correction (SVC; 10-mm radius sphere) was derived from the independent auditory motion localizer contrast EB>SC [motion>static] (Fig.2a right) and was centered on the voxel showing the strongest statistical peak in this contrast (xyz in mm MNI space: −11, −81, −36). ** The coordinate for small volume correction (sphere of 10mm radius) was centred on the Planum Temporale region showing auditory motion selective response in the independent localizer contrast SC [motion>static]. Coordinates are reported in MNI space, and regions surviving whole-brain family-wise error (FWE) correction are identified using the Automated Anatomical Labelling (AAL) atlas. L: Left; R: Right.

| Area | Cluster<br>size (k) | x(mm) | y(mm) | z(mm) | z | t | p(FEW) |
| --- | --- | --- | --- | --- | --- | --- | --- |
| SC [visual motion > static] |  |  |  |  |  |  |  |
| R Lingual Gyrus | 38 | 15 | -78 | -9 | 5.17 | 11.19 | 0.028 |
| R Calcarine Sulcus | 6 | 17 | -86 | 12 | 5.15 | 11.09 | 0.03 |
| EB > SC [auditory motion > static] |  |  |  |  |  |  |  |
| L Cuneus | 271 | -11 | -81 | 36 | 3.17 | 3.82 | 0.033* |
| SC > EB [auditory motion > static] |  |  |  |  |  |  |  |
| R Planum Temporale | 115 | 62 | -33 | 13 | 3.73 | 4.81 | 0.007** |

## Notes

### Competing Interest Statement

The authors have declared no competing interest.

## REFERENCES

Aitken, F., Menelaou, G., Warrington, O., Koolschijn, R. S., Corbin, N., Callaghan, M. F., & Kok, P. (n.d.). Prior expectations evoke stimulus-specific activity in the deep layers of the primary visual cortex. PLOS BIOLOGY, 19.

Ajina, S., & Bridge, H. (2018). Blindsight relies on a functional connection between hMT+ and the lateral geniculate nucleus, not the pulvinar. PLOS Biology, 16(7), e2005769. 10.1371/journal.pbio.2005769

Ajina, S., Kennard, C., Rees, G., & Bridge, H. (2015). Motion area V5/MT+ response to global motion in the absence of V1 resembles early visual cortex. Brain, 138(1), 164–178. 10.1093/brain/awu328

Allen, B., Spiegel, D. P., Thompson, B., Pestilli, F., & Rokers, B. (2015). Altered white matter in early visual pathways of humans with amblyopia. Vision Research, Amblyopia: A Window into Visual Cortex Development and Recovery of Vision, 114, 48–55. 10.1016/j.visres.2014.12.021

Amedi, A., Hofstetter, S., Maidenbaum, S., & Heimler, B. (2017). Task Selectivity as a Comprehensive Principle for Brain Organization. Trends in Cognitive Sciences, 21(5), 307–310. 10.1016/j.tics.2017.03.007

Arcaro, M. J., Pinsk, M. A., & Kastner, S. (2015). The Anatomical and Functional Organization of the Human Visual Pulvinar. Journal of Neuroscience, 35(27), 9848–9871. 10.1523/JNEUROSCI.1575-14.2015

Barilari, M., Koiso, K., Taylor, P., Gulban, O. F., Glen, D., Bandettini, P., collignon, olivier, & Huber, L. (2026). Layer-MAP (Modular Analysis Pipeline) (Version 0.2.0) [Computer software]. Zenodo. 10.5281/zenodo.18937199

Barkat, T. R., Polley, D. B., & Hensch, T. K. (2011). A critical period for auditory thalamocortical connectivity. Nature Neuroscience, 14(9), 1189–1194. 10.1038/nn.2882

Battal, C. (2018). Decoding Auditory Motion Direction And Location In hMT+/V5 And Planum Temporale Of Sighted And Blind Individuals [PhD Thesis]. UNITN.

Battal, C., Rezk, M., Mattioni, S., Vadlamudi, J., & Collignon, O. (2019). Representation of Auditory Motion Directions and Sound Source Locations in the Human Planum Temporale. Journal of Neuroscience, 39(12), 2208–2220. 10.1523/JNEUROSCI.2289-18.2018

Bavelier, D., & Neville, H. J. (2002). Cross-modal plasticity: Where and how? Nature Reviews Neuroscience, 3(6), 443–452. 10.1038/nrn848

Berman, R. A., & Wurtz, R. H. (2011). Signals Conveyed in the Pulvinar Pathway from Superior Colliculus to Cortical Area MT. Journal of Neuroscience, 31(2), 373–384. 10.1523/JNEUROSCI.4738-10.2011

Born, R. T., & Bradley, D. C. (2005). Structure and function of visual area MT. Annual Review of Neuroscience, 28, 157–189. 10.1146/annurev.neuro.26.041002.131052

Brainard, D. H. (1997). The Psychophysics Toolbox. Spatial Vision, 10(4), 433–436. 10.1163/156856897X00357

Bridge, H., Cowey, A., Ragge, N., & Watkins, K. (2009). Imaging studies in congenital anophthalmia reveal preservation of brain architecture in ‘visual’ cortex. Brain, 132(12), 3467–3480. 10.1093/brain/awp279

Bridge, H., Thomas, O., Jbabdi, S., & Cowey, A. (2008). Changes in connectivity after visual cortical brain damage underlie altered visual function. Brain, 131(6), 1433–1444. 10.1093/brain/awn063

Bronchti, G., Heil, P., Sadka, R., Hess, A., Scheich, H., & Wollberg, Z. (2002). Auditory activation of ‘visual’ cortical areas in the blind mole rat (Spalax ehrenbergi). European Journal of Neuroscience, 16(2), 311–329. 10.1046/j.1460-9568.2002.02063.x

Bronchti, G., Heil, P., Scheich, H., & Wollberg, Z. (1989). Auditory pathway and auditory activation of primary visual targets in the blind mole rat (Spalax ehrenbergi): I. 2-deoxyglucose study of subcortical centers. The Journal of Comparative Neurology, 284(2), 253–274. 10.1002/cne.902840209

Büchel, C. (2003). Cortical hierarchy turned on its head. Nature Neuroscience, 6(7), 657–658. 10.1038/nn0703-657

Callaway, E. M. (1998). LOCAL CIRCUITS IN PRIMARY VISUAL CORTEX OF THE MACAQUE MONKEY. Annual Review of Neuroscience, 21(Volume 21, 1998), 47–74. 10.1146/annurev.neuro.21.1.47

Cecchetti, L., Ricciardi, E., Handjaras, G., Kupers, R., Ptito, M., & Pietrini, P. (2016). Congenital blindness affects diencephalic but not mesencephalic structures in the human brain. Brain Structure and Function, 221(3), 1465–1480. 10.1007/s00429-014-0984-5

Chabot, N., Charbonneau, V., Laramée, M.-E., Tremblay, R., Boire, D., & Bronchti, G. (2008). Subcortical auditory input to the primary visual cortex in anophthalmic mice. Neuroscience Letters, 433(2), 129–134. 10.1016/j.neulet.2008.01.003

Chai, Y., Liu, T. T., Marrett, S., Li, L., Khojandi, A., Handwerker, D. A., Alink, A., Muckli, L., & Bandettini, P. A. (2021). Topographical and laminar distribution of audiovisual processing within human planum temporale. Progress in Neurobiology, 102121. 10.1016/j.pneurobio.2021.102121

Chai, Y., Morgan, A. T., Xie, H., Li, L., Huber, L., Bandettini, P. A., & Sutton, B. P. (2024). Unlocking near-whole-brain, layer-specific functional connectivity with 3D VAPER fMRI. Imaging Neuroscience, 2, 1–20. 10.1162/imag_a_00140

Charbonneau, V., Laramée, M.-E., Boucher, V., Bronchti, G., & Boire, D. (2012). Cortical and subcortical projections to primary visual cortex in anophthalmic, enucleated and sighted mice. European Journal of Neuroscience, 36(7), 2949–2963. 10.1111/j.1460-9568.2012.08215.x

Chou, X., Fang, Q., Yan, L., Zhong, W., Peng, B., Li, H., Wei, J., Tao, H. W., & Zhang, L. I. (2020). Contextual and cross-modality modulation of auditory cortical processing through pulvinar mediated suppression. eLife, 9, e54157. 10.7554/eLife.54157

Cohen, L. G., Celnik, P., Pascual-Leone, A., Corwell, B., Faiz, L., Dambrosia, J., Honda, M., Sadato, N., Gerloff, C., Catala, M. D., & Hallett, M. (1997). Functional relevance of cross-modal plasticity in blind humans. Nature, 389(6647), 180–183. 10.1038/38278

Collignon, O., Dormal, G., Albouy, G., Vandewalle, G., Voss, P., Phillips, C., & Lepore, F. (2013). Impact of blindness onset on the functional organization and the connectivity of the occipital cortex. Brain: A Journal of Neurology, 136(Pt 9), 2769–2783. 10.1093/brain/awt176

Collignon, O., Dormal, G., de Heering, A., Lepore, F., Lewis, T. L., & Maurer, D. (2015). Long-Lasting Crossmodal Cortical Reorganization Triggered by Brief Postnatal Visual Deprivation. Current Biology, 25(18), 2379–2383. 10.1016/j.cub.2015.07.036

Collignon, O., Dormal, G., & Lepore, F. (2012). Building the brain in the dark: Functional and specific crossmodal reorganization in the occipital cortex of blind individuals. Plasticity in Sensory Systems, 73, 96.

Collignon, O., Vandewalle, G., Voss, P., Albouy, G., Charbonneau, G., Lassonde, M., & Lepore, F. (2011). Functional specialization for auditory–spatial processing in the occipital cortex of congenitally blind humans. Proceedings of the National Academy of Sciences, 108(11), 4435–4440. 10.1073/pnas.1013928108

Cox, R. W. (1996). AFNI: Software for Analysis and Visualization of Functional Magnetic Resonance Neuroimages. Computers and Biomedical Research, 29(3), 162–173. 10.1006/cbmr.1996.0014

Crair, M. C., & Malenka, R. C. (1995). A critical period for long-term potentiation at thalamocortical synapses. Nature, 375(6529), 325–328. 10.1038/375325a0

Dehay, C., Giroud, P., Berland, M., Killackey, H., & Kennedy, H. (1996). Contribution of thalamic input to the specification of cytoarchitectonic cortical fields in the primate: Effects of bilateral enucleation in the fetal monkey on the boundaries, dimensions, and gyrification of striate and extrastriate cortex. Journal of Comparative Neurology, 367(1), 70–89. 10.1002/(SICI)1096-9861(19960325)367:1%3C70::AID-CNE6%3E3.0.CO;2-G

Desimone, R., & Ungerleider, L. G. (1986). Multiple visual areas in the caudal superior temporal sulcus of the macaque. Journal of Comparative Neurology, 248(2), 164–189. 10.1002/cne.902480203

Dormal, G., Lepore, F., & Collignon, O. (2012). Plasticity of the Dorsal “Spatial” Stream in Visually Deprived Individuals. Neural Plasticity, 2012(1), 687659. 10.1155/2012/687659

Dormal, G., Rezk, M., Yakobov, E., Lepore, F., & Collignon, O. (2016). Auditory motion in the sighted and blind: Early visual deprivation triggers a large-scale imbalance between auditory and “visual” brain regions. NeuroImage, 134, 630–644. 10.1016/j.neuroimage.2016.04.027

Doron, N., & Wollberg, Z. (1994). Cross-modal neuroplasticity in the blind mole rat Spalax Ehrenbergi: A WGA-HRP tracing study. NeuroReport, 5(18), 2697.

Douglas, R. J., & Martin, K. A. (1991). A functional microcircuit for cat visual cortex. The Journal of Physiology, 440, 735–769. 10.1113/jphysiol.1991.sp018733

Dresbach, S., Huber, L. (Renzo), Gulban, O. F., & Goebel, R. (2023). Layer-fMRI VASO with short stimuli and event-related designs at 7 T. NeuroImage, 279, 120293. 10.1016/j.neuroimage.2023.120293

Dumoulin, S. O. (2000). A New Anatomical Landmark for Reliable Identification of Human Area V5/MT: A Quantitative Analysis of Sulcal Patterning. Cerebral Cortex, 10(5), 454–463. 10.1093/cercor/10.5.454

Ewall, G., Parkins, S., Lin, A., Jaoui, Y., & Lee, H.-K. (2021). Cortical and Subcortical Circuits for Cross-Modal Plasticity Induced by Loss of Vision. Frontiers in Neural Circuits, 15. 10.3389/fncir.2021.665009

Feldman, D. E., Nicoll, R. A., Malenka, R. C., & Isaac, J. T. (1998). Long-term depression at thalamocortical synapses in developing rat somatosensory cortex. Neuron, 21(2), 347–357. 10.1016/s0896-6273(00)80544-9

Felleman, D. J., & Van Essen, D. C. (1991). Distributed Hierarchical Processing in the Primate Cerebral Cortex. Cerebral Cortex, 1(1), 1–47. 10.1093/cercor/1.1.1

Finn, E. S., Huber, L., Jangraw, D. C., Molfese, P. J., & Bandettini, P. A. (2019). Layer-dependent activity in human prefrontal cortex during working memory. Nature Neuroscience, 22(10), 1687–1695. 10.1038/s41593-019-0487-z

Fischl, B. (2012). FreeSurfer. NeuroImage, 20 YEARS OF fMRI, 62(2), 774–781. 10.1016/j.neuroimage.2012.01.021

Frasnelli, J., Collignon, O., Voss, P., & Lepore, F. (2011). Crossmodal plasticity in sensory loss. In Progress in Brain Research (Vol. 191, pp. 233–249). Elsevier. 10.1016/B978-0-444-53752-2.00002-3

Gaglianese, A., Costagli, M., Bernardi, G., Ricciardi, E., & Pietrini, P. (2012). Evidence of a direct influence between the thalamus and hMT+ independent of V1 in the human brain as measured by fMRI. NeuroImage, 60(2), 1440–1447. 10.1016/j.neuroimage.2012.01.093

Gau, R., Barilari, M., Battal, C., Rezk, M., Collignon, O., Gurtubay, A., Falagiarda, F., MacLean, M., Cerpelloni, F., Shahzad, I., Nunes, M., Caron-Guyon, J., Chouinard-Leclaire, C., Yang, Y., Mattioni, S., Van Audenhaege, A., & Matuszewski, J. (n.d.). Bidspm (Version 4.0.0) [Computer software]. Retrieved https://github.com/cpp-lln-lab/bidspm

Gau, R., Bazin, P.-L., Trampel, R., Turner, R., & Noppeney, U. (2020). Resolving multisensory and attentional influences across cortical depth in sensory cortices. eLife, 9, e46856. 10.7554/eLife.46856

Gorgolewski, K. J., Auer, T., Calhoun, V. D., Craddock, R. C., Das, S., Duff, E. P., Flandin, G., Ghosh, S. S., Glatard, T., Halchenko, Y. O., Handwerker, D. A., Hanke, M., Keator, D., Li, X., Michael, Z., Maumet, C., Nichols, B. N., Nichols, T. E., Pellman, J., … Poldrack, R. A. (2016). The brain imaging data structure, a format for organizing and describing outputs of neuroimaging experiments. Scientific Data, 3(1), 160044. 10.1038/sdata.2016.44

Huber, L., Benedikt, P., Bandettini, P., Arora, K., Wagstyl, K., Cho, S., Goense, J., Nothnagel, N., Morgan, A. T., van den Hurk, J., Müller, A. K., Reynolds, R., Glen, D., Goebel, R., & Gulban, O. F. (2021). LayNii: A software suite for layer-fMRI (Version v2.3.0) [Computer software]. Zenodo. 10.5281/zenodo.7586233

Huber, L., Finn, E. S., Chai, Y., Goebel, R., Stirnberg, R., Stöcker, T., Marrett, S., Uludag, K., Kim, S.-G., Han, S., Bandettini, P. A., & Poser, B. A. (2021). Layer-dependent functional connectivity methods. Progress in Neurobiology, How High Spatiotemporal Resolution fMRI Can Advance Neuroscience, 207, 101835. 10.1016/j.pneurobio.2020.101835

Huber, L., Goense, J., Kennerley, A. J., Trampel, R., Guidi, M., Reimer, E., Ivanov, D., Neef, N., Gauthier, C. J., Turner, R., & Möller, H. E. (2015). Cortical lamina-dependent blood volume changes in human brain at 7T. NeuroImage, 107, 23–33. 10.1016/j.neuroimage.2014.11.046

Huber, L., Handwerker, D. A., Jangraw, D. C., Chen, G., Hall, A., Stüber, C., Gonzalez-Castillo, J., Ivanov, D., Marrett, S., Guidi, M., Goense, J., Poser, B. A., & Bandettini, P. A. (2017). High-Resolution CBV-fMRI Allows Mapping of Laminar Activity and Connectivity of Cortical Input and Output in Human M1. Neuron, 96(6), 1253–1263.e7. 10.1016/j.neuron.2017.11.005

Huber, L., Ivanov, D., Krieger, S. N., Streicher, M. N., Mildner, T., Poser, B. A., Möller, H. E., & Turner, R. (2014). Slab-selective, BOLD-corrected VASO at 7 Tesla provides measures of cerebral blood volume reactivity with high signal-to-noise ratio. Magnetic Resonance in Medicine, 72(1), 137–148. 10.1002/mrm.24916

Huber, L., Poser, B. A., Bandettini, P. A., Arora, K., Wagstyl, K., Cho, S., Goense, J., Nothnagel, N., Morgan, A. T., Hurk, J. van den, Müller, A. K., Reynolds, R. C., Glen, D. R., Goebel, R., & Gulban, O. F. (2021). LayNii: A software suite for layer-fMRI. NeuroImage, 237, undefined-undefined. 10.1016/j.neuroimage.2021.118091

Huber, L., Poser, B., Bandettini, P., Arora, K., Wagstyl, K., Cho, S., Goense, J., Nothnagel, N., Morgan, A. T., van den Hurk, J., Müller, A. K., Reynolds, R., Glen, D., Goebel, R., & Gulban, O. F. (2024). LayNii: A software suite for layer-fMRI (Version v2.6.0) [Computer software]. Zenodo. 10.5281/zenodo.10523194

Huber, L., Uludağ, K., & Möller, H. E. (2019). Non-BOLD contrast for laminar fMRI in humans: CBF, CBV, and CMRO2. NeuroImage, 197, 742–760. 10.1016/j.neuroimage.2017.07.041

Ibrahim, L. A., Mesik, L., Ji, X.-Y., Fang, Q., Li, H.-F., Li, Y.-T., Zingg, B., Zhang, L. I., & Tao, H. W. (2016). Cross-Modality Sharpening of Visual Cortical Processing through Layer-1-Mediated Inhibition and Disinhibition. Neuron, 89(5), 1031–1045. 10.1016/j.neuron.2016.01.027

Iurilli, G., Ghezzi, D., Olcese, U., Lassi, G., Nazzaro, C., Tonini, R., Tucci, V., Benfenati, F., & Medini, P. (2012). Sound-Driven Synaptic Inhibition in Primary Visual Cortex. Neuron, 73(4–2), 814–828. 10.1016/j.neuron.2011.12.026

Jiang, B., Treviño, M., & Kirkwood, A. (2007). Sequential Development of Long-Term Potentiation and Depression in Different Layers of the Mouse Visual Cortex. Journal of Neuroscience, 27(36), 9648–9652. 10.1523/JNEUROSCI.2655-07.2007

Jiang, F., Stecker, G. C., Boynton, G. M., & Fine, I. (2016). Early Blindness Results in Developmental Plasticity for Auditory Motion Processing within Auditory and Occipital Cortex. Frontiers in Human Neuroscience, 10(July). 10.3389/fnhum.2016.00324

Kahn, D. M., & Krubitzer, L. (2002). Massive cross-modal cortical plasticity and the emergence of a new cortical area in developmentally blind mammals. Proceedings of the National Academy of Sciences of the United States of America, 99(17), 11429–11434. 10.1073/pnas.162342799

Kaiserman-Abramof, I. R., Graybiel, A. M., & Nauta, W. J. H. (1980). The thalamic projection to cortical area 17 in a congenitally anophthalmic mouse strain. Neuroscience, 5(1), 41–52. 10.1016/0306-4522(80)90069-X

Karlen, S. J., Kahn, D. M., & Krubitzer, L. (2006). Early blindness results in abnormal corticocortical and thalamocortical connections. Neuroscience, 142(3), 843–858. 10.1016/j.neuroscience.2006.06.055

Kim, S.-G. (2025). The road to mapping neural circuits with fMRI: Challenges and advances. Aperture Neuro, 5(SI 1). 10.52294/001c.138741

Kleiner, M., Brainard, D., & Pelli, D. (2007). What’s new in Psychtoolbox-3? Perception - ECVP Abstract Supplement. European Conference on Visual Perception (ECVP-2007), August 27-31, Arezzo, Italy. 10.1177/03010066070360S101

Klinge, C., Eippert, F., Röder, B., & Büchel, C. (2010). Corticocortical Connections Mediate Primary Visual Cortex Responses to Auditory Stimulation in the Blind. Journal of Neuroscience, 30(38), 12798–12805. 10.1523/JNEUROSCI.2384-10.2010

Koiso, K., Müller, A. K., Akamatsu, K., Dresbach, S., Wiggins, C. J., Gulban, O. F., Goebel, R., Miyawaki, Y., Poser, B. A., & Huber, L. (2022). Acquisition and processing methods of whole-brain layer-fMRI VASO and BOLD: The Kenshu dataset (p. 2022.08.19.504502). bioRxiv. 10.1101/2022.08.19.504502

Lakatos, P., Chen, C.-M., O’Connell, M. N., Mills, A., & Schroeder, C. E. (2007). Neuronal oscillations and multisensory interaction in primary auditory cortex. Neuron, 53(2), 279–292. 10.1016/j.neuron.2006.12.011

Lankinen, K., Ahlfors, S. P., Mamashli, F., Blazejewska, A. I., Raij, T., Turpin, T., Polimeni, J. R., & Ahveninen, J. (2022). Cortical depth profiles of auditory and visual 7 T functional MRI responses in human superior temporal areas. Human Brain Mapping, n/a(n/a). 10.1002/hbm.26046

Lawrence, S. J. D., Formisano, E., Muckli, L., & de Lange, F. P. (2019). Laminar fMRI: Applications for cognitive neuroscience. NeuroImage, 197, 785–791. 10.1016/j.neuroimage.2017.07.004

Leclerc, C., Saint-Amour, D., Lavoie, M. E., Lassonde, M., & Lepore, F. (2000). Brain functional reorganization in early blind humans revealed by auditory event-related potentials. NeuroReport, 11(3), 545.

Leh, S. E., Chakravarty, M. M., & Ptito, A. (2008). The Connectivity of the Human Pulvinar: A Diffusion Tensor Imaging Tractography Study. International Journal of Biomedical Imaging, 2008(1), 789539. 10.1155/2008/789539

López-Bendito, G., Aníbal-Martínez, M., & Martini, F. J. (2022). Cross-Modal Plasticity in Brains Deprived of Visual Input Before Vision. Annual Review of Neuroscience, 45(Volume 45, 2022), 471–489. 10.1146/annurev-neuro-111020-104222

Macaluso, E., & Driver, J. (2005). Multisensory spatial interactions: A window onto functional integration in the human brain. Trends in Neurosciences, 28(5), 264–271. 10.1016/j.tins.2005.03.008

Macaluso, E., Frith, C. D., & Driver, J. (2000). Modulation of Human Visual Cortex by Crossmodal Spatial Attention. Science, 289(5482), 1206–1208. 10.1126/science.289.5482.1206

Marques, J. P., Kober, T., Krueger, G., van der Zwaag, W., Van de Moortele, P.-F., & Gruetter, R. (2010). MP2RAGE, a self bias-field corrected sequence for improved segmentation and T1-mapping at high field. NeuroImage, 49(2), 1271–1281. 10.1016/j.neuroimage.2009.10.002

Mattioni, S., Rezk, M., Battal, C., Vadlamudi, J., & Collignon, O. (2022). Impact of blindness onset on the representation of sound categories in occipital and temporal cortices. eLife, 11, e79370. 10.7554/eLife.79370

Mattioni, S., Rezk, M., Gao, X., Nam, J., Liu, Z.-X., Gau, R., Goffaux, V., Costantino, A. I., Op de Beeck, H., Lewis, T., Maurer, D., & Collignon, O. (2025). Impact of a transient neonatal visual deprivation on the development of the ventral occipito-temporal cortex in humans. Nature Communications, 16(1), 9828. 10.1038/s41467-025-65468-7

Matuszewski, J., Bola, Ł., Collignon, O., & Marchewka, A. (2025). Similar Computational Hierarchies for Reading and Speech in the Occipital Cortex of Sighed and Blind: Converging Evidence from fMRI and Chronometric TMS. The Journal of Neuroscience, 45(20), e1153242024. 10.1523/JNEUROSCI.1153-24.2024

Maunsell, J. H. R., & Van Essen, D. C. (1983). The connections of the middle temporal visual area (MT) and their relationship to a cortical hierarchy in the macaque monkey. The Journal of Neuroscience, 3(12), 2563–2586. 10.1523/JNEUROSCI.03-12-02563.1983

Merabet, L. B., & Pascual-Leone, A. (2010). Neural reorganization following sensory loss: The opportunity of change. Nature Reviews Neuroscience, 11(1), 44–52. 10.1038/nrn2758

Møller, H., Sørensen, M. F., Jensen, C. B., & Hammershøi, D. (1996). Binaural technique: Do we need individual recordings? Journal of the Audio Engineering Society, 44(6), 451–469.

Movshon, J. A., & Newsome, W. T. (1996). Visual Response Properties of Striate Cortical Neurons Projecting to Area MT in Macaque Monkeys. Journal of Neuroscience, 16(23), 7733–7741. 10.1523/JNEUROSCI.16-23-07733.1996

Muckli, L., De Martino, F., Vizioli, L., Petro, L. S., Smith, F. W., Ugurbil, K., Goebel, R., & Yacoub, E. (2015). Contextual Feedback to Superficial Layers of V1. Current Biology, 25(20), 2690–2695. 10.1016/j.cub.2015.08.057

Müller, F., Niso, G., Samiee, S., Ptito, M., Baillet, S., & Kupers, R. (2019). A thalamocortical pathway for fast rerouting of tactile information to occipital cortex in congenital blindness. Nature Communications, 10(1), 5154. 10.1038/s41467-019-13173-7

Oude Lohuis, M. N., Marchesi, P., Olcese, U., & Pennartz, C. M. A. (2024). Triple dissociation of visual, auditory and motor processing in mouse primary visual cortex. Nature Neuroscience, 27(4), 758–771. 10.1038/s41593-023-01564-5

Pelli, D. G. (1997). The VideoToolbox software for visual psychophysics: Transforming numbers into movies. Spatial Vision, 10(4), 437–442.

Poirier, C., Collignon, O., Scheiber, C., Renier, L., Vanlierde, A., Tranduy, D., Veraart, C., & De Volder, A. G. (2006). Auditory motion perception activates visual motion areas in early blind subjects. NeuroImage, 31(1), 279–285. 10.1016/j.neuroimage.2005.11.036

Ponce, C. R., Lomber, S. G., & Born, R. T. (2008). Integrating motion and depth via parallel pathways. Nature Neuroscience, 11(2), 216–223. 10.1038/nn2039

Ptito, M., Schneider, F. C. G., Paulson, O. B., & Kupers, R. (2008). Alterations of the visual pathways in congenital blindness. Experimental Brain Research, 187(1), 41–49. 10.1007/s00221-008-1273-4

R Core Team. (2024). R: A Language and Environment for Statistical Computing. R Foundation for Statistical Computing. https://www.R-project.org/

Rezk, M., Cattoir, S., Battal, C., Occelli, V., Mattioni, S., & Collignon, O. (2020). Shared Representation of Visual and Auditory Motion Directions in the Human Middle-Temporal Cortex. Current Biology, S0960982220305534. 10.1016/j.cub.2020.04.039

Ricciardi, E., Vanello, N., Sani, L., Gentili, C., Scilingo, E. P., Landini, L., Guazzelli, M., Bicchi, A., Haxby, J. V., & Pietrini, P. (2007). The Effect of Visual Experience on the Development of Functional Architecture in hMT+. Cerebral Cortex, 17(12), 2933–2939. 10.1093/cercor/bhm018

Rockland, K. S. (1989). Bistratified distribution of terminal arbors of individual axons projecting from area V1 to middle temporal area (MT) in the macaque monkey. Visual Neuroscience, 3(2), 155–170. 10.1017/S0952523800004466

Rockland, K. S. (1995). Morphology of individual axons projecting from area V2 to MT in the macaque. Journal of Comparative Neurology, 355(1), 15–26. 10.1002/cne.903550105

Rons, T. (1996). Novel sensations in the congenitally blind. Nature, 380(6574), 479–480. 10.1038/380479a0

Sadato, N., Pascual-Leone, A., Grafman, J., Ibañez, V., Deiber, M.-P., Dold, G., & Hallett, M. (1996). Activation of the primary visual cortex by Braille reading in blind subjects. Nature, 380(6574), 526–528. 10.1038/380526a0

Schroeder, C. E., & Foxe, J. J. (2002). The timing and laminar profile of converging inputs to multisensory areas of the macaque neocortex. Cognitive Brain Research, Multisensory Proceedings, 14(1), 187–198. 10.1016/S0926-6410(02)00073-3

Stirnberg, R., & Stöcker, T. (2021). Segmented K-space blipped-controlled aliasing in parallel imaging for high spatiotemporal resolution EPI. Magnetic Resonance in Medicine, 85(3), 1540–1551. 10.1002/mrm.28486

Tustison, N. J., Cook, P. A., Holbrook, A. J., Johnson, H. J., Muschelli, J., Devenyi, G. A., Duda, J. T., Das, S. R., Cullen, N. C., Gillen, D. L., Yassa, M. A., Stone, J. R., Gee, J. C., & Avants, B. B. (2021). The ANTsX ecosystem for quantitative biological and medical imaging. Scientific Reports, 11(1), 9068. 10.1038/s41598-021-87564-6

Vizioli, L., Moeller, S., Dowdle, L., Akçakaya, M., De Martino, F., Yacoub, E., & Uğurbil, K. (2021). Lowering the thermal noise barrier in functional brain mapping with magnetic resonance imaging. Nature Communications, 12(1), 5181. 10.1038/s41467-021-25431-8

Watson, J. D. G., Myers, R., Frackowiak, R. S. J., Hajnal, J. V., Woods, R. P., Mazziotta, J. C., Shipp, S., & Zeki, S. (1993). Area V5 of the Human Brain: Evidence from a Combined Study Using Positron Emission Tomography and Magnetic Resonance Imaging. Cerebral Cortex, 3(2), 79–94. 10.1093/cercor/3.2.79

Yaka, R., Yinon, U., Rosner, M., & Wollberg, Z. (2000). Pathological and experimentally induced blindness induces auditory activity in the cat primary visual cortex. Experimental Brain Research, 131(1), 144–148. 10.1007/s002219900295

Zeki, S. (1991). CEREBRAL AKINETOPSIA (VISUAL MOTION BLINDNESS): A REVIEW. Brain, 114(2), 811–824. 10.1093/brain/114.2.811

## REFERENCES

Barilari, M., Koiso, K., Taylor, P., Gulban, O. F., Glen, D., Bandettini, P., Collignon, Olivier, & Huber, L. (2026). Layer-MAP (Modular Analysis Pipeline) (Version 0.2.0) [Computer software]. Zenodo. 10.5281/zenodo.18937199

Huber, L., Benedikt, P., Bandettini, P., Arora, K., Wagstyl, K., Cho, S., Goense, J., Nothnagel, N., Morgan, A. T., van den Hurk, J., Müller, A. K., Reynolds, R., Glen, D., Goebel, R., & Gulban, O. F. (2024). LayNii: A software suite for layer-fMRI (Version v2.6.0) [Computer software]. Zenodo. 10.5281/zenodo.10523194

Huber, L. (Renzo), Poser, B. A., Bandettini, P. A., Arora, K., Wagstyl, K., Cho, S., Goense, J., Nothnagel, N., Morgan, A. T., van den Hurk, J., Müller, A. K., Reynolds, R. C., Glen, D. R., Goebel, R., & Gulban, O. F. (2021). LayNii: A software suite for layer-fMRI. NeuroImage, 237, 118091. 10.1016/j.neuroimage.2021.11809154

